# Frontal Theta Oscillations and Cognitive Flexibility: Age-Related Modulations in EEG Activity

**DOI:** 10.1101/2024.07.05.602082

**Authors:** Margarita Darna, Christopher Stolz, Hannah-Sophia Jauch, Hendrik Strumpf, Jens-Max Hopf, Constanze I. Seidenbecher, Björn H. Schott, Anni Richter

## Abstract

Cognitive flexibility, the ability to adapt one’s behaviour in changing environments, declines during aging. Electroencephalography (EEG) studies have implicated midfrontal theta oscillations in attentional set-shifting, a measure of cognitive flexibility. Little is known about the electrocortical underpinnings of set-shifting in aging. Here, we investigated aging effects on set-shifting performance by analysing theta power in 20 young (mean age: 22.5 ± 2.9 years) and 19 older (mean age: 69.4 ± 6.1 years) adults. Increasing shift difficulty (i.e., intra- vs. extra-dimensional shifts) elicited worse performance in both age groups, with older adults showing overall longer reaction times (RTs) and increased RT variability. Young adults exhibited amplified midfrontal theta power increases with higher shift difficulty whereas older adults showed overall lower theta power and no task-related midfrontal theta power modulation, indicating potentially distinct underlying neural mechanisms.

## 1. Introduction

Cognitive flexibility refers to the ability to adapt one’s behaviour in a changing environment. As an executive function (Diamond, 2013), cognitive flexibility relies on more basic cognitive functions, such as attention, selection processes and performance monitoring (Cavanagh and Frank, 2014). Cognitive flexibility is commonly investigated using task-switching and set-shifting paradigms, which require to flexibly adapt one’s response to repeatedly changing stimulus-response mappings. Such paradigms typically elicit switch costs, namely increased reaction times (RTs) and error rates when a rule is changed compared to trials (or blocks) without change (Wylie and Allport, 2000). Many set-shifting studies have distinguished between intra-dimensional (ID) and extra-dimensional (ED) shifts. During an ID shift, features change within a dimension, whereas in ED shifts, the rule is changed to a different feature dimension. ED shifts are likely more difficult than ID shifts and thus engage more neural resources (Watson et al., 2006).

Studies on changes in cognitive flexibility in older age yet report inconsistent findings. While some report deficits (Reynolds et al., 1995; Richter et al., 2023; Wecker et al., 2005), mostly evident in increased switch costs (Cepeda et al., 2001; Kray et al., 2002; Meiran et al., 2001), more perseverative errors (Haaland et al., 1987; Zelazo et al., 2004), and failure to perform ED shifts (De Luca et al., 2003; Owen et al., 1991; Zelazo et al., 2004), others document that switch costs are unaltered (Falkenstein et al., 2001; Karayanidis et al., 2011; Kolev et al., 2005; Kray and Lindenberger, 2000) or even diminished (Kray, 2006) in older adults. Apart from these inconsistent behavioural results during set-shifting in older adults, little is known about their neural underpinnings.

Task-related analyses of electroencephalography (EEG) data, such as event-related time-frequency analysis, can help to decipher the neural mechanisms underlying alterations of cognitive flexibility in old age. For the present study, we focused on midfrontal theta oscillations (4-8 Hz) a key EEG parameter previously discussed in the context of cognitive flexibility. In general, midfrontal theta is relevant in the context of cognitive control processes (Cavanagh and Shackman, 2015; Domic-Siede et al., 2021; Stolz et al., 2023), action adjustment (van de Vijver et al., 2011), maintenance of working memory content (Ratcliffe et al., 2022), and cognitive interference (Nigbur et al., 2011). Its source has been localized in the anterior cingulate cortex and adjacent medial prefrontal cortex (Cavanagh and Shackman, 2015; Tsujimoto et al., 2006), a core structure of the brain’s salience network (Seeley, 2019). Regarding cognitive flexibility, switch trials have been associated with increased theta power compared to repeat trials (Cooper et al., 2017; Cunillera et al., 2012) and increased theta phase-coupling between prefrontal and posterior (i.e., occipital and temporo-parietal) regions (Sauseng et al., 2006).

In older individuals, there are conflicting results regarding theta and cognitive flexibility. Lower theta power during baseline activity and lower theta coherence across brain regions during set-shifting was observed (Dias et al., 2015), as well as a lack of frontal theta modulation during reversal learning (Küçük et al., 2023). Conversely, theta power increase in frontal regions was found during attentional switching in older age, whereas youngest participants exhibited parietal theta modulations (Huizeling et al., 2021). After separated analysis of ID and ED set-shifting conditions to acknowledge their proposed different needs for cognitive resources, Oh et al. (2014) observed young adults’ theta activity in the inferior frontal gyrus to be higher in ED compared to ID trials, yet this finding has to be proved in older adults.

To this end, we applied the EEG-compatible ID/ED set-shifting (IDED) task (Oh et al., 2014) in which we assessed whether set-shifting deficits in healthy older adults might be reflected by the modulation of theta by shift types (ID vs. ED). We expected midfrontal theta power to be overall decreased in older participants and increased in ED as compared to ID shifts in young individuals, reflecting the need for higher cognitive control with increasing set-shifting demands. In older adults we expected theta modulation in set-shifting to differ from young adults, either being reduced or absent. Finally, we hypothesized a relationship between midfrontal theta power and individual differences in set-shifting performance.

## 2. Materials and Methods

### 2.1 Participants

Participants were between 18 and 35 years old or at least 60 years old to be assigned to the young or the older age groups, respectively. All participants gave written informed consent in accordance with the Declaration of Helsinki (World Medical Association, 2013) and received financial compensation for participation. The study was approved by the Ethics Committee of the Faculty of Medicine at the Otto von Guericke University of Magdeburg. Exclusion criteria for participation were manifested neurological or psychiatric disorders or history thereof, past or present substance dependence or abuse, use of neurological or psychiatric medication, and serious medical conditions (e.g., heart failure NYHA stage III or IV, metastatic cancer, or diabetes mellitus with complications) as assessed by a routine neuropsychiatric interview and a health questionnaire. All participants were right-handed and had fluent German language skills. Crystallized intelligence was estimated with the “Mehrfachwahl-Wortschatz-Intelligenztest B” (MWT-B) (Lehrl et al., 1995).

Older participants also completed the Mini Mental State Examination (MMSE: Folstein et al., 1975) to evaluate their cognitive status. None of the participants scored below 26, and thus well above the level of 24, which has been suggested as a value indicative for dementia in a comprehensive meta-analysis (e.g. Creavin et al., 2016).

The experiment was performed by a total of 41 participants, 20 young (mean age: 22.5 ± 2.9 years) and 21 older. This sample size was chosen based on previous research (e.g. Oh et al., 2014; Yeung et al., 2016) and according to the minimum sample size required (*N* = 34) to achieve 80% power for a medium effect size (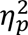= 0.05) at a significance criterion of *α* = .05 for a within-between interaction in an analysis of variance (ANOVA) (G*Power version 3.1.9.1; Faul et al., 2009). Two older participants were excluded from further analysis due to high noise in the EEG signal resulting from muscle artifacts, or high error rates (>50%) resulting in 19 older participants included in the final study sample (mean age: 69.4 ± 6.1 years; Table 1).

**Table 1:**
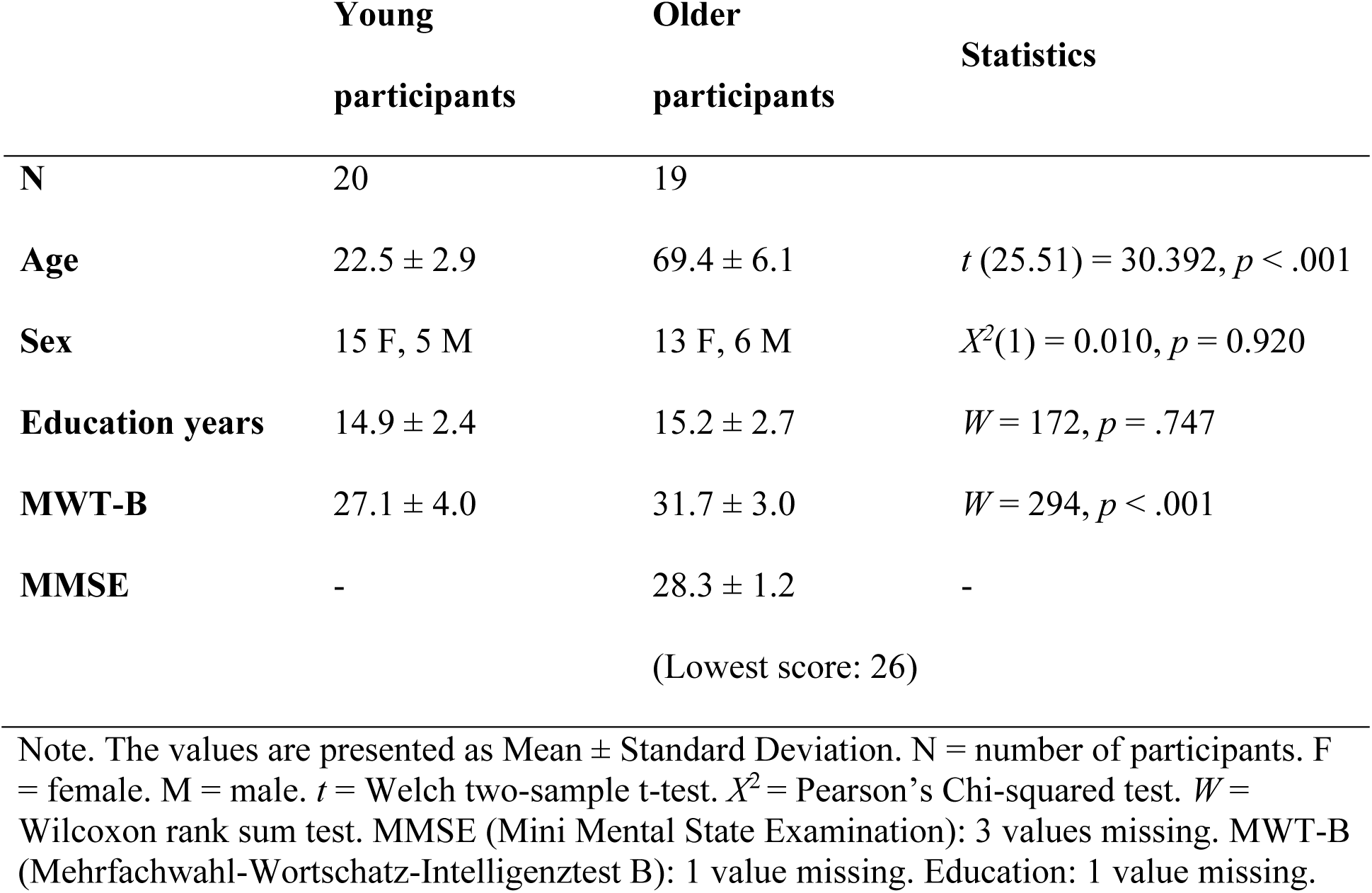
Participant summary statistics.

|  | Young<br>participants | Older<br>participants | Statistics |
| --- | --- | --- | --- |
| **N** | 20 | 19 |  |
| **Age** | 22.5 ± 2.9 | 69.4 ± 6.1 | $t(25.51) = 30.392, p < .001$ |
| **Sex** | 15 F, 5 M | 13 F, 6 M | $X^2(1) = 0.010, p = 0.920$ |
| **Education years** | 14.9 ± 2.4 | 15.2 ± 2.7 | $W = 172, p = .747$ |
| **MWT-B** | 27.1 ± 4.0 | 31.7 ± 3.0 | $W = 294, p < .001$ |
| **MMSE** | - | 28.3 ± 1.2 | - |
|  |  | (Lowest score: 26) |  |

Note. The values are presented as Mean ± Standard Deviation. N = number of participants. F = female. M = male. $t$ = Welch two-sample t-test. $X^2$ = Pearson's Chi-squared test. $W$ = Wilcoxon rank sum test. MMSE (Mini Mental State Examination): 3 values missing. MWT-B (Mehrfachwahl-Wortschatz-Intelligenztest B): 1 value missing. Education: 1 value missing.

### 2.2 Cognitive Flexibility Task

To assess participants’ cognitive flexibility, all participants first completed the Attentional Set Shifting Task (ASST; Sahakian and Owen, 1992). The results and a brief discussion are provided in the supplementary material (Supplementary Material S3).

In order to investigate the set-shifting ability with EEG, we used the IDED: an adapted version of the ASST, previously established and used with MEG (Oh et al., 2014).

The IDED is a trial-based paradigm in which participants match one of two stimuli presented to a single target. During each trial, only one of the two stimuli matches the target in one of two possible dimensions: colour or shape. The matching rule stays the same for at least three consecutive trials, during which the target also remains unchanged. When a new target appears a change of the matching rule occurs, initiating an ID or ED set shift. Within ID shifts, the dimension remains the same but a different feature value needs to be attended (colour-to-colour or shape-to-shape) (Figure 1C) whereas in ED trials, the previously irrelevant dimension becomes relevant (colour-to-shape or shape-to-colour) (Figure 1D).

**Figure 1:**
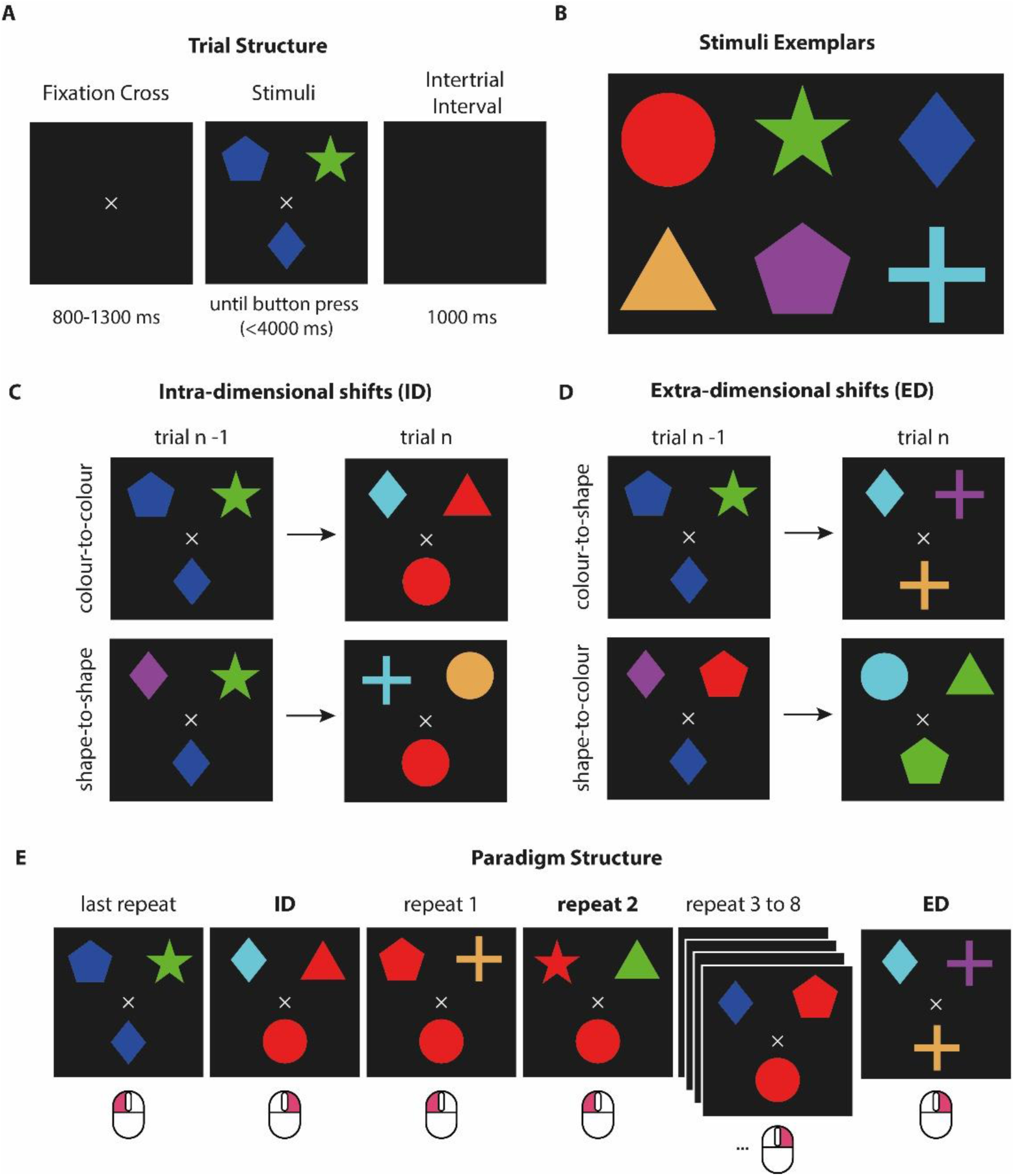
**The IDED Task**. **A:** Trial structure. After the presentation of the fixation cross for 800 to 1300 ms the stimuli and target appear. The participants signal with a righthand mouse click which stimulus matches the target either by colour or shape (here: left button press to indicate colour match). After button press or after a time of 4000 ms without response the stimuli and target disappear. An empty screen is visible for 1000 ms followed by a new trial; **B:** Stimuli exemplars showcasing the six possible colour (red, green, blue, beige, magenta, cyan) and shape combinations (circle, star, diamond, triangle, pentagon, cross) resulting in 36 possible stimuli; **C:** Types of intra-dimensional shifts (ID). ID shifts could either be colour-to-colour (e.g. blue-to-red, top row) or shape-to-shape (e.g. diamond-to-circle, bottom row); **D:** Types of extra-dimensional shifts (ED). ED shifts could either be colour-to-shape (e.g. blue-to-cross, top row) or shape-to-colour (e.g. diamond-to-green, bottom row); **E:** Paradigm structure. Here, an example of possible consequent trials is presented. In each trial the target matches one of the stimuli by colour or by shape. In ID or ED trials the target changes its form and colour eliciting a shift, followed by a pseudorandomized number of repeat trials (2 to 7 trials), in which both the target and matching rule remain the same. Below each panel the correct mouse response (left or right) is visible. Trials in bold represent the conditions of interest.

Each trial always had the following structure (Figure 1A). Within each trial, a fixation cross was presented on a black background (Figure 1A). After a pseudorandomized interval of 800 to 1300 ms (generated with uniformly distributed pseudorandom numbers in MATLAB; The MathWorks Inc., 2021; https://www.mathworks.com; Version 9.11.0.1809720; R2021b), the stimuli and target were presented above and below the fixation cross, respectively. The participant had a total of 4 seconds to respond via button press. Pressing the left versus right button meant that the left versus right stimulus matched the target in its colour or shape, respectively. If no button was pressed after four seconds the trial was terminated and counted as an error trial. After button press or trial termination, a black screen was presented for 1000 ms, followed by the fixation cross, which signalled the beginning of the next trial.

Six possible colours (red, green, blue, beige, magenta and cyan) and six possible shapes (circle, star, diamond, triangle, pentagon and cross) were combined to create 36 distinct stimuli (Figure 1B). Before the start of the experiment, all participants were shown the six colours in the form of squares to ensure that they could successfully differentiate the six colours from each other. Participants then completed a set of 20 practice trials, in which all conditions were included and feedback was given, as part of task preparation. The practice trials were not included in the analysis. After practice, no further feedback on response correctness was given.

Participants were not aware in advance which dimension was relevant in each trial. In total 50 ED (25 colour-to-shape and 25 shape-to-colour) and 50 ID (25 colour-to-colour and 25 shape-to-shape) trials were presented. Shift trials were followed by 2 to 7 trials in which the target and matching rule remained the same (e.g., in case of a colour match the target remained a red square and one of the stimuli was red). Trials were presented in a pseudorandomized order that was established by shuffling the type of shifts (ID or ED) and the number of repeat trials after them (2 to 7) in MATLAB. The Toolbox Psychtoolbox was used for stimulus presentation (Brainard and Vision, 1997; http://psychtoolbox.org/; Version 3.0.18) and port event signalling was performed with the Mex-File Plug-in IO64 (http://apps.usd.edu/coglab/psyc770/IO64.html).

Only the second trial after a shift was used as a control trial in the analysis (Oh et al., 2014). These trials will be referred to as repeat trials. The participants completed the task in 6 blocks with 5 self-paced breaks. After a break all trials that occurred before the first shift were excluded from the analysis. The task duration was approximately 30 to 40 min.

Recordings took place in a dimmed electrically shielded room. Stimuli were presented on a 32” presentation monitor with a refresh rate of 120 Hz and a resolution of 1920 x 1280 pixels. The participants were seated at a distance of 110 cm away from the monitor.

Responses were recorded as mouse clicks with the right hand.

### 2.3 EEG Acquisition and Pre-processing

EEG was recorded at a sampling rate of 1000 Hz from 64 active electrodes (Brain Products GmbH, Gilching, Germany) of the actiCAP layout (EASYCAP) and additional reference and ground electrodes (online reference: FCz; ground: AFz) using BrainRecorder from Brain Products. Electrode impedance was kept below 10 kΩ with electrolyte gel.

Data were pre-processed using Brain Vision Analyzer (Brain Products GmbH, Gilching, Germany). The continuous EEG data of the remaining electrodes were first re-referenced to the average of the TP9 and TP10 channels, representing virtually linked mastoids. The data were then filtered with zero phase shift Butterworth filters and a 1 Hz to 100 Hz band-pass second-order filter (Winkler et al., 2015). Artefacts were detected semi-automatically by defining subject-specific criteria based on the gradient (mean maximum allowed voltage step: 50 *μV*, marked time window: 400 ms before and after an event), maximum difference of values in an interval length of 200 ms (mean maximal allowed absolute difference: 600 *μV*, marked time window: 1000 ms before and after an event) and low activity in intervals of 100 ms (lowest allowed activity: 0.5 *μV*, marked time window: 200 ms before and after an event; Supplementary Table 1). We opted for individual thresholds for the gradient criterion as this accounts for individual variability in the EEG signal and differences in noise level (Luck, 2014). Here, channels with more than 20% rejected data points were interpolated using spline interpolation (order 2, degree 10 and *λ =* 1*e^-5^*). Channel interpolation was conducted in four participants, for one channel each (Supplementary Table 2). Next, Independent Component Analysis (ICA) was performed to detect components of blinks and saccades. ICA weight matrices were saved and applied to the data with the same pre-processing steps as described above but using different filters: A 0.1 to 100 Hz band-pass second order filter was applied for wavelet analysis. The previously detected blink and saccade ICA components were removed by a trained observer (author MD). A second artefact detection was performed here to ensure the detection of artefacts that may have been missed prior to the ICA (the maximum difference of values in an interval length of 200 ms was set to 400 *μV*; the other criteria remained the same). The continuous data sets were then down-sampled to 500 Hz and exported for further processing in MATLAB using Fieldtrip (Oostenveld et al., 2011; http://fieldtriptoolbox.org; Version 20211122).

In Fieldtrip, custom scripts were used for wavelet analysis including the following steps: a) extraction of stimulus-locked epochs (from -1250 ms until 3000 ms) from the trials of interest (repeat, ID, and ED trials) that were followed by correct responses (a lengthy stimulus-locked epoch was chosen to account for the relatively low frequency of theta oscillations, as at least 7 full theta cycles were required for the wavelet analysis and to enable a better investigation of response trials), b) baseline correction using the average of -250 to 0 ms pre-stimulus.

Frequency power from 1 to 60 Hz in steps of 1 Hz was then calculated using Morlet wavelets with a width of 7 cycles and length equal to 3 standard deviations of the implicit Gaussian kernel. The time window used was sliding in steps of 50 ms, where 0 was defined as the onset of stimulus presentation. A baseline of -400 ms to -200 ms of the common average of each participant was used for normalization of all trials within each frequency bin separately to avoid temporal smearing (Morales and Bowers, 2022), as done in previous studies (Panitz et al., 2019). Normalization into *dB* was performed using the following equation:

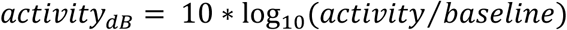

Midfrontal theta power was calculated as the mean theta power (4-8 Hz) at electrode sites F1, F2, Fz, FC1, FC2, FCz in the post-stimulus time period between 250 and 500 ms. These electrodes and time period were chosen based on the observed distribution of theta activity on the grand average of all participants and all conditions (Stolz et al., 2023). The same time window was chosen for both age groups, as the onset of theta power increase after stimulus presentation did not differ between the two age groups (see supplementary material S2). These locations also correspond to the positions used for analysis of midfrontal theta power in other studies, in particular the FCz (e.g. Kolev et al., 2024; Ratcliffe et al., 2022; Yordanova et al., 2020).

For the purposes of single-trial analysis, the mean theta power obtained from the frontocentral electrodes in the time period between 250 and 500 ms after target presentation, was calculated for each trial before normalization with baseline subtraction. Single-trial correlations were calculated separately for each participant and each condition using Shepherd’s Pi correlation (Schwarzkopf et al., 2012), which enables unbiased outlier removal by bootstrapping based on the Mahalanobis distance. The obtained correlation coefficients were then transformed in z-space using Fisher’ z-transformation to make them approach normal distribution for further statistical analysis, resulting in single trial z-scores.

### 2.4 Statistical Analysis

All statistical analyses were performed in RStudio (R Core Team, 2022; R version 4.2.2; R Studio Team, 2020).

IDED reaction time (RT) was defined as the time taken to correctly respond after stimulus presentation. Participants’ RTs exhibited a right-skewed distribution (mean skewness of all trials: 2.30 ± 0.75) and were therefore first log-transformed (log_RTs). The mean and standard deviation of the distribution of the logarithmic RTs (SD_log_RT) were used as dependent variables in statistical analyses. Error rates were computed as percentage of trials with incorrect or no responses in each condition.

Performance z-scores were computed as the additive inverse of the standardized scores for each performance measure (log_RTs, standard deviation of log_RTs and error rate), according to:

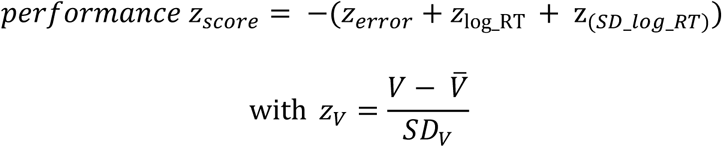

where *V* denotes the individual score on a given performance measure and *V̄* and 𝑆𝐷_*V*_ represent the mean and standard deviation of that measure across all participants and conditions (Dias et al., 2015; Liesefeld and Janczyk, 2019). *V̄* and 𝑆𝐷_*V*_ were weighted to account for unequal sample sizes across age groups and performance z-scores are a negative sum of the individual scores, ensuring that higher performance z-scores indicate better performance (i.e., fewer errors, faster responses, and less variability in response times).

Alpha significance level was set to *p* = .05. ANOVA results are reported with the obtained *F* value, *p* value and partial eta squared (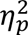) as a measure of effect size. Whenever ANOVAs were significant, post-hoc two-tailed t-tests were performed to uncover specific differences between the means. Cohen’s d is reported as effect size. Statistical significance is reported in the graphs as followed: * for 𝑝 ≤ .05 , ** for 𝑝 ≤ .005 and *** for 𝑝 ≤ .001. In graphs the error bars represent the 95% confidence interval around the mean unless otherwise stated. Average values in text are reported with standard deviations in the form: mean ± standard deviation, unless otherwise specified.

The following variables were tested as dependent variables in separate two-factorial mixed ANOVAs with the between-subjects factor age group (young vs. older adults) and the within-subjects factor condition (repeat vs. ID vs. ED): mean log_RT, SD_log_RT, error rate, performance z-scores, midfrontal theta power, and the single trial z-scores. To investigate the spatial specificity of the observed theta effects, we performed a three-factorial mixed ANOVA with central midline electrode position as additional within-subject factor (Fz, FCz, Cz, CPz, Pz, POz and Oz). Sphericity for ANOVAs was tested using Mauchly’s Test. When sphericity assumptions were not met, degrees of freedom were adjusted using the Greenhouse-Geisser correction. All dependent variables were tested for non-normality and inhomogeneity of variances with the Shapiro-Wilk test and Levene’s test respectively. Both tests were non-significant across all variables, conditions, and age groups (Shapiro-Wilk: all *p* > 0.10; Levene: all *p* > 0.05) except for the error rate. Here, non-normality was given (highest value: ED within young participants: *W* = 0.85, *p* = 0.005). Therefore, error rates were additionally compared with a robust ANOVA. The results of the robust ANOVA did not qualitatively differ from the results of the ANOVA and are reported in the supplementary material (S4).

The single trial z-scores were tested against zero using one-sample t-tests. These were corrected for multiple comparisons with the False Discovery Rate (FDR; Benjamini and Hochberg, 1995).

To evaluate the relationship between theta power and performance, we employed a linear mixed effect model on channels FCz and Oz. The model included the main effects and interactions of theta power, age group and condition. To account for subject-level variability a random slope and random intercept for theta power was included for each participant. To assess the significance of the interaction of the fixed effect, we implemented an ANOVA on the fitted linear mixed-effect model using Satterthwaite’s approximation for degrees of freedom. We additionally extracted estimated fixed-effect slopes from the model and used one-sample t-tests to determine whether the slopes differed across conditions and age groups. We also assessed differences in slopes between conditions and age groups using independent-samples t-tests and corrected p-values using FDR.

Whenever accepting the null hypothesis – rather than merely not rejecting it – was appropriate for the interpretation of our results, we employed Bayesian statistics using JASP ( version 0.18.3; JASP Team, 2024). Specifically, we computed Bayesian repeated measures ANOVAs of mean midfrontal theta power within each age group for the factor condition followed by Bayesian paired t-tests. We additionally performed Bayesian one sample t-tests of the single-trial correlation coefficients.

## 3. Results

### 3.1 Behavioural Results of the IDED Task

Across the three conditions, all participants had an overall median RT of 718 ± 199 ms. Mean log RTs were significantly longer in older compared to young participants in all conditions (*F* (1, 37) = 62.00, *p* < .001, 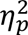= 0.63) (Figure 2A). Regardless of age group, a significant condition effect on log_RTs was found (*F* (1.30, 47.94) = 163.57, *p* < .001, 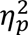= 0.82). As expected, we observed the longest log_RTs in the ED condition, followed by the ID condition, and shortest log_RTs in the repeat condition (ED vs repeat: *t(38)* = 14.80, *p* < .001, *d* = 2.37; ED vs ID: *t(38)* = 9.27, *p* < .001, *d* = 1.48; ID vs repeat: *t(38)* = 15.1, *p* < .001, *d* = 2.43). The interaction between condition and age group did not reach significance (*p =* .727).

**Figure 2:**
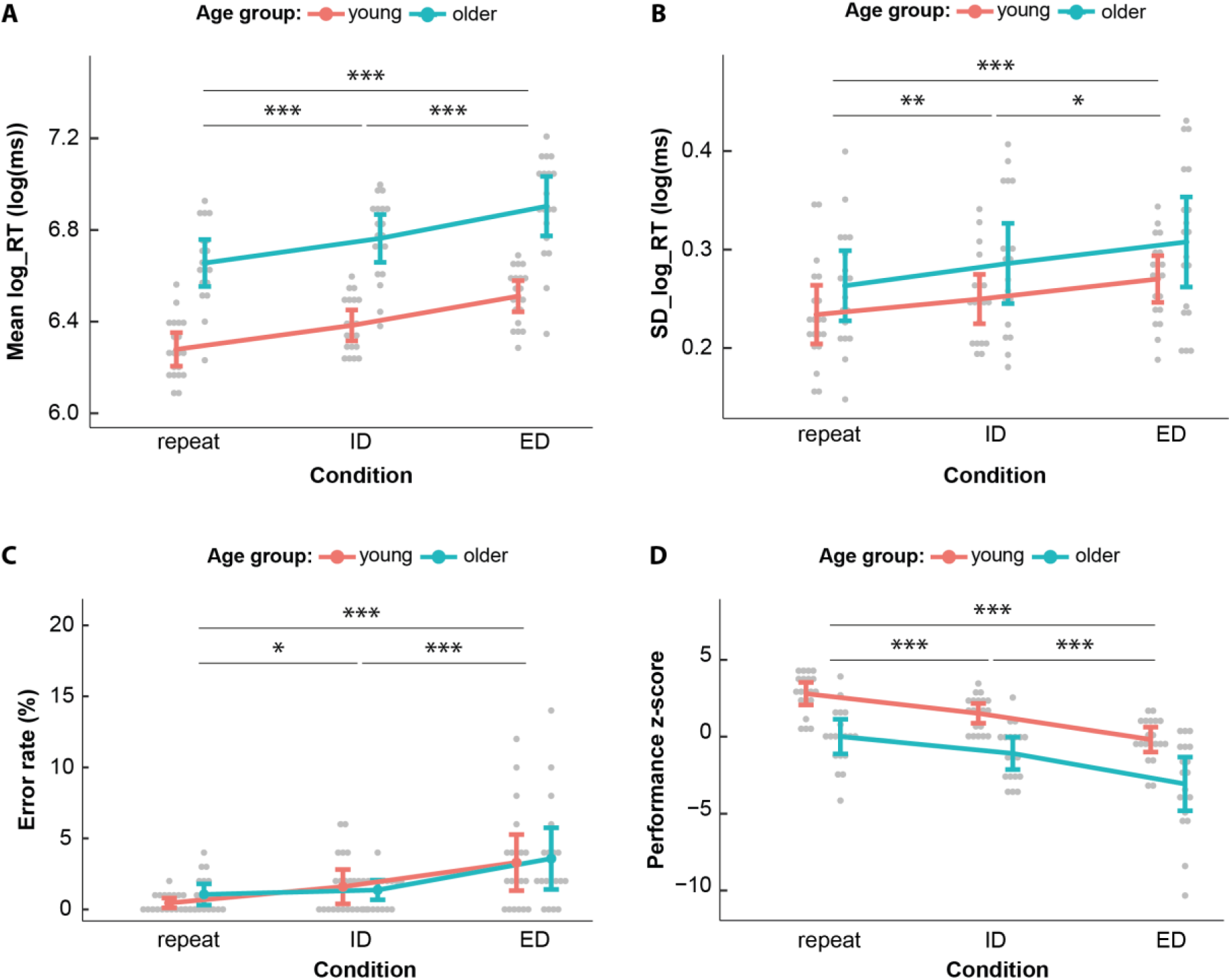
Behavioural results of the IDED task. A: Mean log transformed reaction times (log_RT). There was a significant difference in log_RTs between all conditions in both age groups with ED trials reaching the longest log_RTs followed by ID and repeat (*F* (1.3, 47.94) = 163.57, *p* < .001, 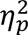= 0.82). Across conditions, older participants had longer log_RTs than younger participants (*F* (1, 37) = 62.00, *p* < .001, 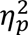= 0.63) (significance not depicted); **B:** Standard deviation of log_RTs (SD_log_RT). There was a significant effect of condition (*F* (2, 74) = 11.49, *p* < .001, 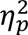= 0.237) and a significant effect of age group (*F* (1, 37) = 4.61, *p* = .038, 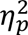= 0.111) (significance not depicted); **C:** Individual error rates. In both age groups there is a gradual increase in error rate from repeat to ID and ED (F (1.32, 48.91) = 17.27, p < .001, 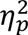= 0.32); **D:** Performance z-scores. There was a significant effect of condition (*F* (1.56, 57.86) = 70.94, *p <* .001, 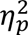= 0.66) and a significant effect of age group (*F* (1, 37) = 29.51, *p <* .001, 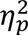= 0.44) (significance not depicted).

In a mixed ANOVA with the SD_log_RTs, we found a significant main effect of condition (*F* (2, 74) = 11.49, *p* < .001, 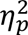= 0.237). Overall, the ED condition resulted in the highest SD_log_RTs, followed by the ID and the repeat condition (ED vs repeat: *t*(38) = 4.89, *p* < .001, *d* = 0.78; ED vs ID: *t* (38) = 2.12, *p* = .040, *d* = 0.34; ID vs repeat: *t*(38) = 2.98, *p* = .005, *d* = 0.48). Additionally, a main effect of age group was found (*F* (1, 37) = 4.61, *p* = .038, 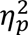= 0.11), where older participants had overall higher SD_log_RTs compared to the younger participants (Figure 2B).

Average error rates were low (1.90% ± 2.60%, see Figure 2C). There was neither a significant difference in error rates between the age groups (*p* = .68), nor a significant interaction of age group and condition (*p* = .59), but a significant main effect of condition was found (*F* (1.32, 48.91) = 17.27, *p <* .001, 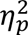= 0.32; Figure 2C). Again, the worst performance with most errors was observed in the ED trials followed by ID trials, and best performance in the repeat trials (ED vs repeat: *t(38)* = 5.07, *p*< .001, *d* = 0.81; ED vs ID: *t(38)* = 3.50, *p* = .001, *d* = 0.56; ID vs repeat: *t(38)* = 2.87, *p* = .007, *d* = 0.46).

Analyses on the composite performance z-scores revealed a significant effect of age group (*F* (1, 37) = 29.51, *p <* .001, 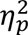= 0.44), with older adults having an overall worse performance. Additionally, a main effect of condition was found (*F* (1.56, 57.86) = 70.94, *p <* .001, 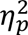= 0.66), where the ED trials displayed the lowest performance z-scores, followed by ID trials and with repeat trials exhibiting the highest z-scores (ED vs repeat: *t(38)* = 10.80, *p*< .001, *d* = 1.72; ED vs ID: *t(38)* = 6.39, *p* = .001, *d* = 1.02; ID vs repeat: *t(38)* = 6.82, *p* = .007, *d* = 1.09). No significant interaction was found (*p =* .796) (Figure 2D).

### 3.2 Midfrontal Theta Activity

During the post-stimulus interval, theta oscillations showed the expected timing and topography in the group average, as compared to a previous study (Oh et al., 2014). Theta first appeared over posterior occipital electrodes between 0 and 200 ms (Figure 3), which can be attributed to the visual evoked response after stimulus presentation, followed by midfrontal activations in the area of the electrodes F1, F2, Fz, FC1, FC2 and FCz. Midfrontal theta power gradually increased from 250 to 500 ms and reached a maximum around 300 ms after stimulus presentation (Figure 4A&B).

**Figure 3:**
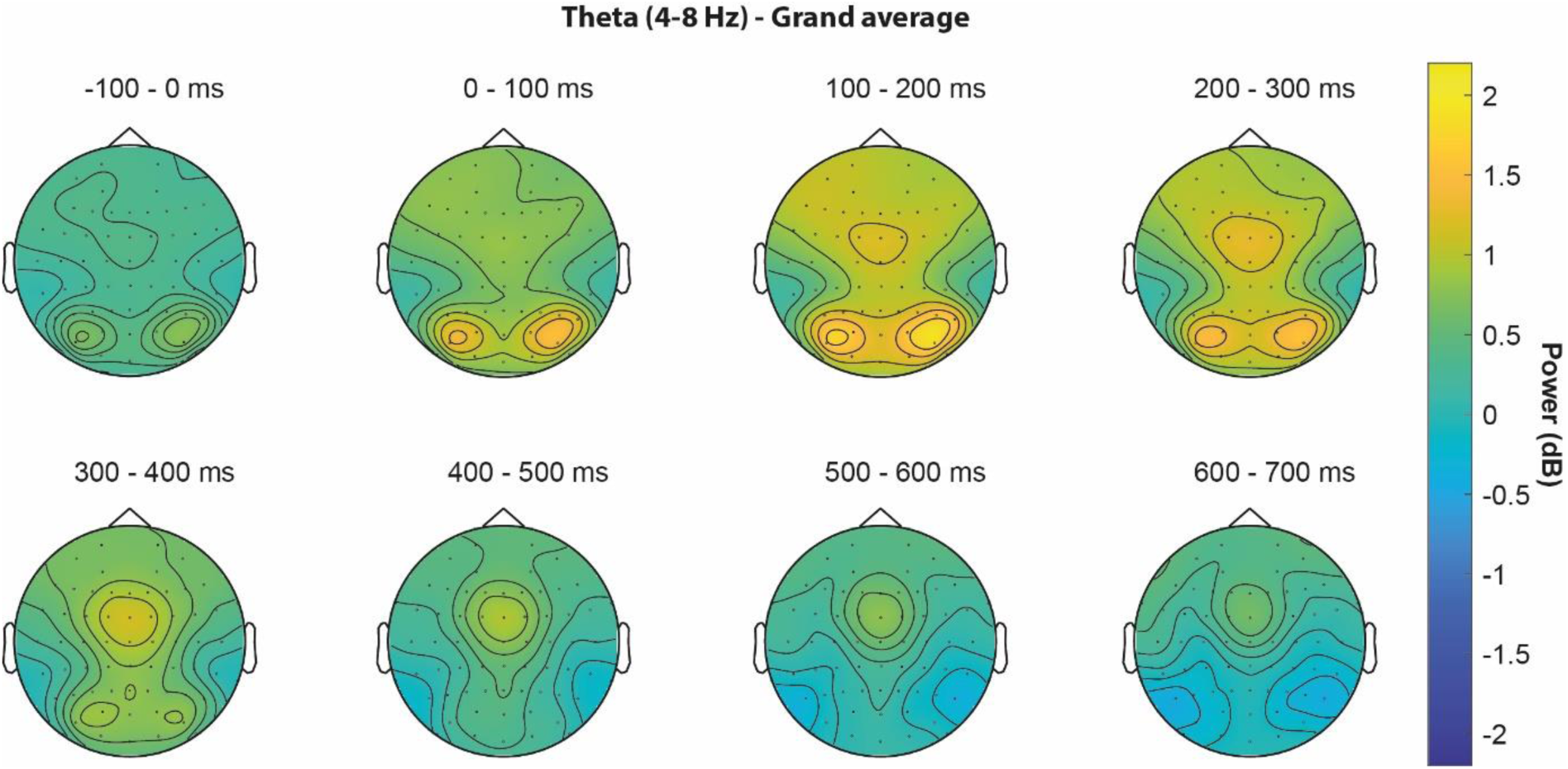
**Mean theta power (dB) across all conditions and all participants**. Occipital theta activations after stimulus presentation (0 to 200 ms) are followed by frontocentral theta activations in the time window between 200 to 500 ms after target presentation.

**Figure 4:**
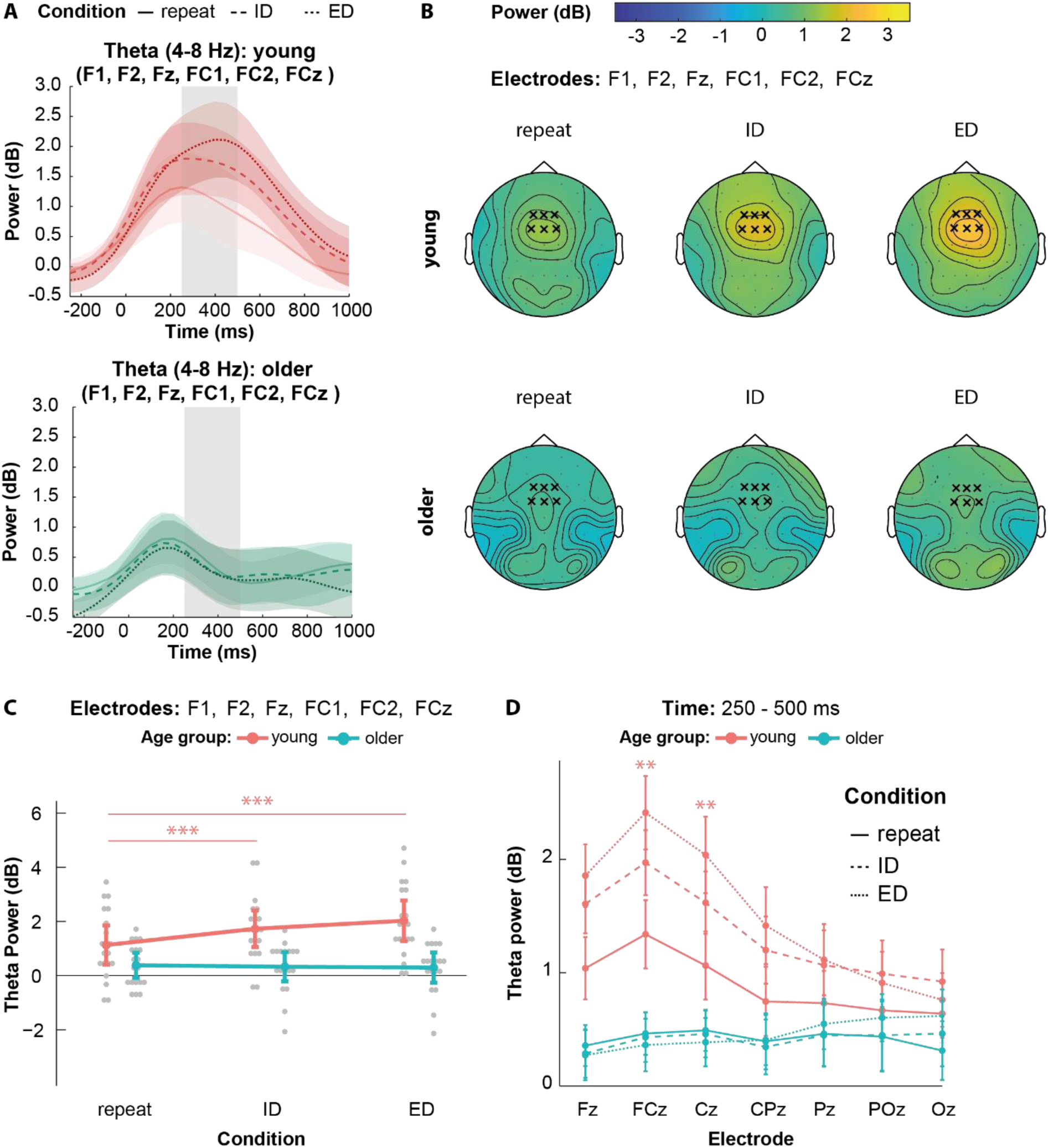
Condition-dependent modulation of midfrontal theta power. A: Average midfrontal theta power after target presentation in the young (top) and older (bottom) age groups. The grey shaded area indicates the time period used to calculate mean theta power (250 to 500 ms); **B:** Average topographical distribution of mean theta power in the young (top) and old (bottom) age groups 250-500 ms after target presentation for each condition. The frontocentral electrodes F1, F2, Fz, FC1, FC2 and FCz used for analysis are marked with an X; **C:** Average theta power at the aforementioned midfrontal electrodes. Here, only young individuals show a significant increase in theta power upon set-shifting compared to the repeat condition (interaction term of age group and condition F (2, 74) = 14.51, p < .001, 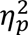= 0.28); **D:** Theta power across the midline electrodes 250-500 ms after target presentation. The asterisks indicate the electrodes in which young participants showcase significant difference across all conditions (highest t-value: FCz; repeat vs ED: *t (37)* = 8.70, *p_adj_* < .001, *d* = 0.92). Older participants show no difference in theta power across conditions in any electrode (highest t-value: CPz; repeat vs ED: *t (37)* = 5.05, *p_adj_* < .001, *d* = 0.57).

Mean midfrontal theta power in the interval between 250 and 500 ms after stimulus presentation differed significantly between age groups (*F* (1, 37) = 15.18, *p* < .001, 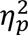= 0.29, *BF_10_* = 52.482), with 1.63 *dB* ± 1.29 *dB* in young vs. 0.34 *dB* ± 0.87 *dB* in older individuals. The latter values did not differ significantly from zero (*p* = .091, *BF_10_*= 0.893), in contrast to the young adults (*t (19)* = 6.04, *p* < .001, *d* = 1.35, BF_10_ = 2.6×10^3^ ). We also observed significant differences in theta power across conditions within the young group, which were largely absent in the older participants (age group by condition interaction: *F* (2, 74) = 14.51, *p* < .001, 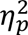= 0.28, *BF_10_* =5.19×10^6^ ) (Figure 4A, B & C). Specifically, in young adults, ED and ID trials elicited an average theta power of 2.03 *dB* ± 1.31 *dB* and 1.73 *dB* ± 1.18 *dB*, respectively. Both the ED and ID conditions elicited significantly higher theta power than the repeat condition, which reached an average of 1.13 *dB* ± 1.26 *dB* (ED vs repeat: *t(19)* = 9.46, *p* < .001, *d* = 2.11, *BF_10_* =9.94×10^5^ ; ID vs repeat: *t(19)* = 4.24, *p* < .001, *d* = 0.95, *BF_10_* =74.72; ED vs ID: *t(19)* = 2.06, *p =* .053, *d* = 0.46, *BF_10_* = 1.33) (Figure 4C). In older individuals, we found no significant differences between conditions (*p* ≥ .780, *BF_10_* ≤ 0.278).

To further evaluate the location specificity of the observed theta effects, we performed a three-way mixed ANOVA with electrode position as additional factor, using the central midline electrodes. Here we found a significant 3-way interaction (*F* (3.47, 128.24) = 6.01, *p* < .001, 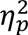= 0.14) (Figure 4D). Post-hoc tests revealed that young adults exhibited significant theta power differences across all conditions at the FCz and Cz electrodes, with the ED condition reaching the highest theta power values, followed by ID and repeat (highest t-value: FCz; repeat vs ED: *t (37)* = 8.70, *p_adj_* < .001, *d* = 0.92). A significant difference in theta power between repeat and one or both of the set-shifting conditions was additionally found in young adults at Fz, CPz and Pz (highest t-value: CPz; repeat vs ED: *t (37)* = 5.05, *p_adj_* < .001, *d* = 0.57). Older adults exhibited no significant differences across conditions in any of the midline electrodes (*p_adj_ ≥* .122) (for detailed results see Supplementary Material S5).

### 3.3 Theta Activity and performance

We next evaluated single trial z-scores of individual RTs and corresponding single-trial theta power at the chosen frontocentral electrodes, using a two-factorial mixed ANOVA. A main effect of age group was found (*F*(1, 37) = 13.73, *p* < .001, 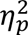= 0.27, *BF_10_* = 18.653), where young participants had higher single trial z-scores than older individuals, reaching values of 0.103 ± 0.176 and -0.025 ± 0.173, respectively. To investigate whether the mean single trial z-score of each age group and condition significantly differed from zero we performed one-sample t-tests of values obtained from each condition and each age group. The z-scores for both the ED and ID conditions among the young individuals significantly differed from zero (ED: *t*(19) = 2.70, 𝑝_𝑎𝑑𝑗_ = .043, *d* = 0.60, *BF_10_* = 3.799; ID *t*(19) = 3.31, 𝑝_𝑎𝑑𝑗_ = .022, *d* = 0.74, *BF_10_* = 11.845)(Figure 5A). In the older adults, none of the values obtained at any condition significantly differed from zero (𝑝_𝑎𝑑𝑗_ ≥ .078, *BF_10_* ≤ 1.722).

**Figure 5:**
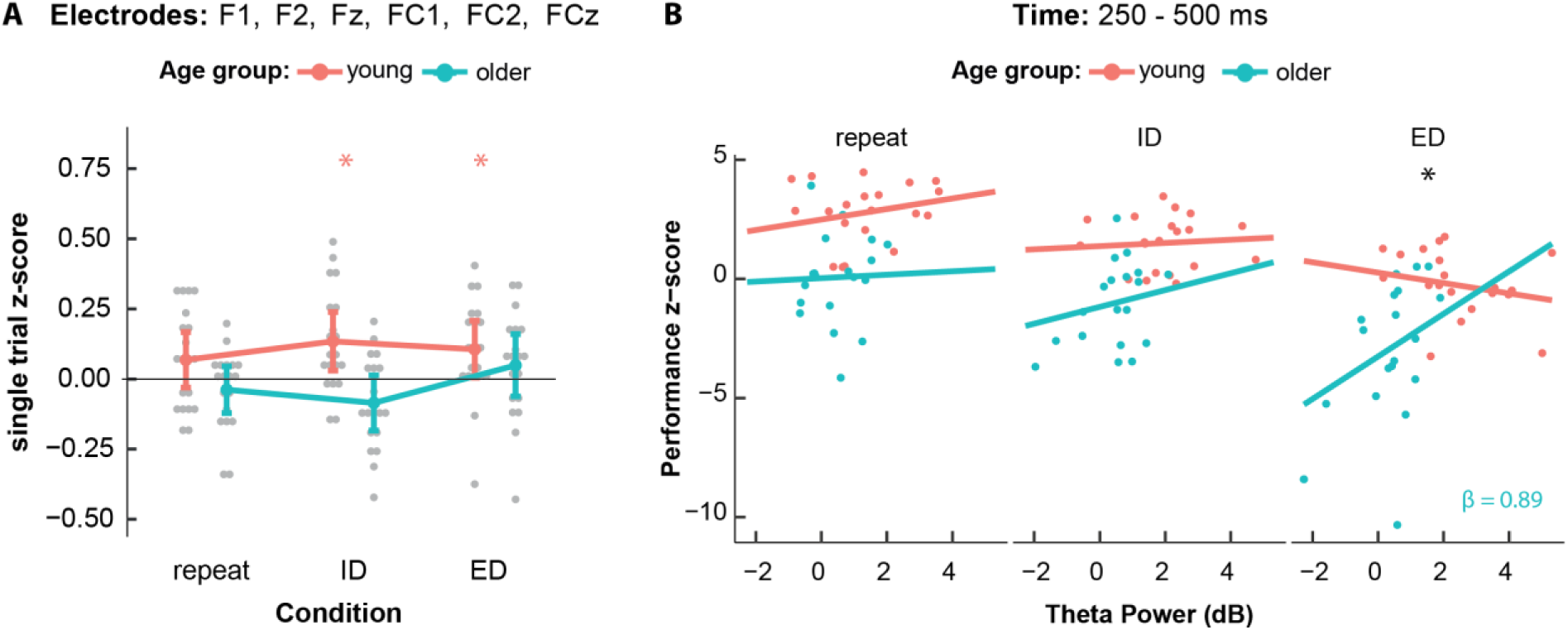
Theta activity and performance A: Single trial z-scores obtained from single trial correlations of theta power and reaction time. Young participants have higher z-values overall (F (1, 37) = 13.73, p < .001, 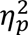= 0.27). Within the young individuals both the ED and ID conditions significantly differ from zero in the positive direction (ED: *t* (19) = 2.70, 𝑝_𝑎𝑑𝑗_ = .043, *d* = 0.60; ID *t* (19) = 3.31, 𝑝_𝑎𝑑𝑗_ = .022, *d* = 0.74); **B:** Estimated slopes of relationship between mean theta power and performance z-scores. In the ED condition, older adults have a significant positive slope which is significantly different from the slope obtained from young participants (*estimate =* 1.11, *SE* = 0.48, *t (71.5)* = 2.30, *p* = .024).

To examine the relationship between performance z-scores and theta power, we employed a linear mixed effects model with theta power on channel FCz, age group and condition as predictor variables for the performance z-scores (for detailed results see Supplementary Material S6). There were significant main effects of age group (*F* (1, 34.53) = 13.95, *p* < .001, 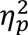= 0.29 ) and condition (*F* (2, 71.59) = 31.27, *p* < .001, 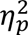= 0.47), mirroring the results that were observed on performance z-scores in 3.1. Additionally, a significant interaction between theta power, age group and condition was found (*F* (2, 72.24) = 3.83, *p =* .026, 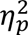= 0.10 ). To investigate this result, we estimated slopes of the relationship between theta power and performance z-scores. Solely the slope of the ED condition in the older age group was significantly different from zero (*SE* = 0.42, *t (83.7)* = 2.13, *p* = .036) reaching a positive value (*β = 0*.89). By comparing the slopes across the groups for each condition, we found that the slope within the ED condition was more positive in older than younger participants (*b =* 1.11, *SE* = 0.48, *t (71.5)* = 2.30, *p* = .024)(Figure 5B).

To assess the spatial specificity of the effects, we also employed a linear mixed effect model with the same specifications at channel Oz, as it was expected that the occipital areas were not involved due to the involvement of the occipital cortex in visual processing Here we found no significant interaction of the three factors (*p =* .059) and none of the estimated slopes were significantly different from zero (*p* ≥ .073).

## 4. Discussion

We observed age-related differences in performance and neural processing during set-shifting. In the IDED task, performance z-scores decreased with higher set-shifting demands and with age. Most notably, older and young adults showed substantial differences in their theta power modulation during the task that we discuss in the following.

### 4.1 Switch costs and age-related changes in cognitive flexibility

Compatible with previous investigations (for a review, see Monsell, 2003), switch costs (ED > ID > repeat) manifested as increasing error rates and longer reaction times, as well as larger log_RT variability in young and older adults. Older adults showed overall lower performance z-scores – considering errors, log_RTs and log_RT variability –, and the majority of older participants failed to complete the ASST, mostly at the ED stage (see S2).

Difficulties in extradimensional shifts have previously been observed in older adults (De Luca et al., 2003; Zelazo et al., 2004). A decline in cognitive flexibility with normal aging has also been shown in task switching (Cepeda et al., 2001; Kray and Lindenberger, 2000) and in other tasks that require cognitive flexibility (Ridderinkhof et al., 2002).

In the – easier – IDED task, there was no conclusive evidence for increased behavioural switch costs in older adults, as both age groups showed equivalent decreases in performance z-scores during set-shifting. Karayanidis et al. (2011) have argued that increased switch costs might not reflect actual set-shifting deficits, but rather increased mixing costs (Kray, 2006; Meiran et al., 2001) which result from using blocks with multiple different trial types. In their specific task, proactive preparation eliminated switch costs. According to the authors, increased switch costs in other task-switching paradigms might reflect insufficient preparation and slowing of non-decision processes like target encoding, suppression of irrelevant information, or slower post-decision processes. It should be noted, though, that their study, like other task-switching studies, relied more on proactive preparation upon cue presentation, whereas our paradigm largely relied on reactive processes. We can thus not exclude that increased switch costs would have become apparent in the IDED, if the participants had less time to employ alternative strategies (see 4.3 Limitations).

### 4.2 Age-related differences in midfrontal theta modulations and cognitive control in set-shifting

In line with the findings in tasks that demand cognitive flexibility (Cooper et al., 2017; Cunillera et al., 2012; Dias et al., 2015) we found an increase in midfrontal theta power during shifts compared to repeat trials in the young group, aligning with the heightened demand for cognitive control during shifting. This modulation was notably observable across the frontocentral electrodes, peaking on FCz as observed previously (Kolev et al., 2024; Ratcliffe et al., 2022; Yordanova et al., 2020). Remarkably, single-trial performance within the young group correlated with theta power, with average z-values within both set-shifting conditions being consistently above zero, indicating a positive correlation between midfrontal theta power and RTs in individual trials. This observation is consistent with similar findings in previous studies on response conflict (Cohen and Cavanagh, 2011; Cohen and van Gaal, 2014).

Our study aligns with previous studies that identified midfrontal theta-band activity as a core neural signature of executive function (Cohen, 2011a, 2014; Duprez et al., 2020). During set-shifting, in particular, midfrontal theta may play a role in updating stimulus-response mappings (van de Vijver et al., 2014), selective attention (McDermott et al., 2017) behaviour re-adjustment (Fusco et al., 2018) and inhibition (Kaiser et al., 2019). Midfrontal theta may also coordinate the interaction between the executive system in frontal areas and the parietally located representational system, as previously reported for working memory (Ratcliffe et al., 2022).

In the older age group, only low midfrontal theta activity was observed, in line with previous studies of cognitive flexibility (Dias et al., 2015), short term memory (Cummins and Finnigan, 2007), and working memory (Gajewski and Falkenstein, 2014). Collectively, these findings provoke the question why older adults exhibited consistently lower midfrontal theta power during task execution, which did not significantly increase with increasing demands on cognitive flexibility.

One possible explanation for this effect could be the functional reorganization of frontal networks with aging (Koen and Rugg, 2019; Reuter-Lorenz, 2002). This has been indirectly observed in recent studies on midfrontal theta where a disengagement of the medial frontal lobe during error processing (Kolev et al., 2024) and during motor coordination (Yordanova et al., 2020) in older age was conveyed. Some past studies have additionally reported a redistribution of medial activity to more frontal areas (van de Vijver et al., 2014; Velanova et al., 2006).

Collectively, a functional rearrangement of frontal cortical function, indicated by altered theta activity, may play a role in diminished cognitive flexibility in older age. One possible explanation for such reorganization could be cortical volume loss as it has been associated with increased response latencies in task switching (Hakun et al., 2015). Prefrontal white matter integrity has also been associated with cognitive control (Ziegler et al., 2010) and memory performance (Cohen, 2011b; Kennedy and Raz, 2009; Schott et al., 2011). Notably, prefrontal white matter tract integrity has also been directly linked to theta amplitude during error processing (Cohen, 2011a). Hence, alterations in theta oscillations in older adults may, at least in part, be attributable to the frequently observed prefrontal cortical volume loss and reduced white matter integrity in aging (Anders M. Fjell et al., 2009; Jernigan et al., 2001 but see: He et al., 2021; Moretti et al., 2007) and/or decreased metabolic activity of the anterior cingulate cortex in older adults (Pardo et al., 2007).

As of note, the positive relationship between mean theta power and performance z-scores in older adults during the ED condition highlights two key points important for future research: First, older participants who showed increased theta power during ED shifts, reaching similar levels to younger participants, also exhibited better performance. This highlights the heterogeneity of cognitive aging with considerable interindividual variability of executive function. Second, the occurrence of the effect in the composite z-score, implicate a more global system of executive function rather than one focused solely on reaction times. This may reflect a focus on reducing errors or maintaining consistent performance across trials – features commonly seen in older age groups as part of the speed-accuracy trade-off (Starns and Ratcliff, 2010).

### 4.3 Limitations of the study

A limitation of our current study is a lack of time-pressure, as participants had a four-second time window to respond in each trial and might therefore not have responded as quickly as possible. This is also reflected by the unexpectedly low error rates observed. These factors may have potentially concealed the increased switch costs expected in older participants. In the study by Dias et al. (2015), WCST performance was assessed differently by examining completed categories and the rates of perseverative and non-perseverative errors. In the IDED, floor effects resulting from low error rates precluded this type of evaluation. We assume that, if we had employed a narrower time window for reactions, we might have observed overall faster response times and a higher frequency of errors, making the heightened switch costs in the older population more conspicuous. Nevertheless, it should be mentioned here that increased switch costs in older age are not invariably observed in studies of task switching (Falkenstein et al., 2001; Kolev et al., 2005).

Finally, we acknowledge the limitations of the small sample size and the limited scope of cognitive assessment in the older participants. We cannot exclude that some older participants may have had subtle cognitive deficits that escaped our cognitive screening (please see supplementary discussion, S.3.3). More studies in the future with larger sample sizes and deeper subject phenotyping shall help better interpret the results and answer questions that arose from this investigation.

### 4.5 Conclusions and future perspectives

In summary, our study highlights the role of midfrontal theta oscillations in age-related differences in set-shifting. Our findings suggest that midfrontal theta oscillatory activity is not necessary for successful set-shifting in older age, indicating that different neural substrates might be recruited for set-shifting due to functional reorganization of frontal networks. Future research utilizing simultaneous EEG-fMRI or MEG to delineate functional connectivity changes with subcortical areas may help to further elucidate the underlying mechanisms.

Maintaining cognitive flexibility in older age is essential, particularly in an aging society. Previous studies show that task switching can be trained (Karbach and Kray, 2009; Strobach et al., 2012), even in older individuals (Cepeda et al., 2001; Steyvers et al., 2019; Whitson et al., 2014). Understanding the underlying neural mechanisms may help to develop more targeted training and intervention protocols. Our results suggest that, in this context, EEG oscillations in particular, might constitute a useful readout, potentially not only in normal aging but also in mild cognitive impairment and Alzheimer’s disease (Yener et al., 2022).

## Acknowledgments

The authors are grateful to Alexander Dityatev for valuable comments on the manuscript. The authors would like to thank Matthias Deliano and Renate Blobel-Lüer for their expert technical assistance. Finally, we would like to express our sincere gratitude to our study participants for their consent to take part in this research study.

## Author Contribution

MD: conceptualization, data curation, methodology, software, formal analysis, investigation, writing – original draft, writing – review and editing, visualization

CS: methodology, writing – original draft, writing – review and editing

HSJ: investigation

HS: methodology

JMH: methodology, writing – original draft, writing – review and editing

BHS: conceptualization, supervision, funding acquisition, writing – original draft, writing – review and editing

CIS: conceptualization, supervision, funding acquisition, writing – review and editing

AR: conceptualization, supervision, methodology, writing – original draft, writing – review and editing

### Data Availability Statement

Experimental Code, MATLAB Scripts, RStudio Scripts and averages of dependent variables are provided in a GitHub repository (https://github.com/margdarna/IDED).

## References

Anders M. Fjell, Kristine B. Walhovd, Christine Fennema-Notestine, Linda K. McEvoy, Donald J. Hagler, Dominic Holland, James B. Brewer, Anders M. Dale, 2009. One-Year Brain Atrophy Evident in Healthy Aging. J. Neurosci. 29, 15223–15231. 10.1523/jneurosci.3252-09.2009

Benjamini, Y., Hochberg, Y., 1995. Controlling the False Discovery Rate: A Practical and Powerful Approach to Multiple Testing. J. R. Stat. Soc. Ser. B Methodol. 57, 289–300. 10.1111/j.2517-6161.1995.tb02031.x

Brainard, D.H., Vision, S., 1997. The psychophysics toolbox. Spat. Vis. 10, 433–436. https://psycnet.apa.org/doi/10.1163/156856897X00357

Cavanagh, J.F., Frank, M.J., 2014. Frontal theta as a mechanism for cognitive control. Trends Cogn. Sci. 18, 414–421. 10.1016/j.tics.2014.04.012

Cavanagh, J.F., Shackman, A.J., 2015. Frontal midline theta reflects anxiety and cognitive control: Meta-analytic evidence. J. Physiol.-Paris 109, 3–15. 10.1016/j.jphysparis.2014.04.003

Cepeda, N.J., Kramer, A.F., Gonzalez de Sather, J.C.M., 2001. Changes in executive control across the life span: Examination of task-switching performance. Dev. Psychol. 37, 715–730. https://psycnet.apa.org/doi/10.1037/0012-1649.37.5.715

Cohen, M.X., 2014. A neural microcircuit for cognitive conflict detection and signaling. Trends Neurosci. 37, 480–490. 10.1016/j.tins.2014.06.004

Cohen, M.X., 2011a. Error-related medial frontal theta activity predicts cingulate-related structural connectivity. NeuroImage 55, 1373–1383. 10.1016/j.neuroimage.2010.12.072

Cohen, M.X., 2011b. Hippocampal-prefrontal connectivity predicts midfrontal oscillations and long-term memory performance. Curr Biol 21, 1900–5. 10.1016/j.cub.2011.09.036

Cohen, M.X., Cavanagh, J.F., 2011. Single-Trial Regression Elucidates the Role of Prefrontal Theta Oscillations in Response Conflict. Front. Psychol. 2. 10.3389/fpsyg.2011.00030

Cohen, M.X., van Gaal, S., 2014. Subthreshold muscle twitches dissociate oscillatory neural signatures of conflicts from errors. NeuroImage 86, 503–513. 10.1016/j.neuroimage.2013.10.033

Cooper, P.S., Wong, A.S.W., McKewen, M., Michie, P.T., Karayanidis, F., 2017. Frontoparietal theta oscillations during proactive control are associated with goal-updating and reduced behavioral variability. Biol. Psychol. 129, 253–264. 10.1016/j.biopsycho.2017.09.008

Creavin, S.T., Wisniewski, S., Noel-Storr, A.H., Trevelyan, C.M., Hampton, T., Rayment, D., Thom, V.M., Nash, K.J.E., Elhamoui, H., Milligan, R., et al, 2016. Mini-Mental State Examination (MMSE) for the detection of dementia in clinically unevaluated people aged 65 and over in community and primary care populations. Cochrane Database Syst. Rev. 10.1002/14651858.cd011145.pub2

Cummins, T.D.R., Finnigan, S., 2007. Theta power is reduced in healthy cognitive aging. Int. J. Psychophysiol. 66, 10–17. 10.1016/j.ijpsycho.2007.05.008

Cunillera, T., Fuentemilla, L., Periañez, J., Marco-Pallarès, J., Krämer, U.M., Càmara, E., Münte, T.F., Rodríguez-Fornells, A., 2012. Brain oscillatory activity associated with task switching and feedback processing. Cogn. Affect. Behav. Neurosci. 12, 16–33. 10.3758/s13415-011-0075-5

De Luca, C.R., Wood, S.J., Anderson, V., Buchanan, J.-A., Proffitt, T.M., Mahony, K., Pantelis, C., 2003. Normative Data From the Cantab. I: Development of Executive Function Over the Lifespan. J. Clin. Exp. Neuropsychol. 25, 242–254. 10.1076/jcen.25.2.242.13639

Diamond, A., 2013. Executive functions. Annu. Rev. Psychol. 64, 135–168. 10.1146/annurev-psych-113011-143750

Dias, N.S., Ferreira, D., Reis, J., Jacinto, L.R., Fernandes, L., Pinho, F., Festa, J., Pereira, M., Afonso, N., Santos, N.C., Cerqueira, J.J., Sousa, N., 2015. Age effects on EEG correlates of the Wisconsin Card Sorting Test. Physiol. Rep. 3, e12390. 10.14814/phy2.12390

Domic-Siede, M., Irani, M., Valdés, J., Perrone-Bertolotti, M., Ossandón, T., 2021. Theta activity from frontopolar cortex, mid-cingulate cortex and anterior cingulate cortex shows different roles in cognitive planning performance. NeuroImage 226, 117557. 10.1016/j.neuroimage.2020.117557

Duprez, J., Gulbinaite, R., Cohen, M.X., 2020. Midfrontal theta phase coordinates behaviorally relevant brain computations during cognitive control. NeuroImage 207, 116340. 10.1016/j.neuroimage.2019.116340

Falkenstein, M., Hoormann, J., Hohnsbein, J., 2001. Changes of error-related ERPs with age. Exp. Brain Res. 138, 258–262. 10.1007/s002210100712

Faul, F., Erdfelder, E., Buchner, A., Lang, A.-G., 2009. Statistical power analyses using G* Power 3.1: Tests for correlation and regression analyses. Behav. Res. Methods 41, 1149–1160. 10.3758/BRM.41.4.1149

Folstein, M.F., Folstein, S.E., McHugh, P.R., 1975. “Mini-mental state”: A practical method for grading the cognitive state of patients for the clinician. J. Psychiatr. Res. 12, 189–198. 10.1016/0022-3956(75)90026-6

Fusco, G., Scandola, M., Feurra, M., Pavone, E.F., Rossi, S., Aglioti, S.M., 2018. Midfrontal theta transcranial alternating current stimulation modulates behavioural adjustment after error execution. Eur. J. Neurosci. 48, 3159–3170. 10.1111/ejn.14174

Gajewski, P.D., Falkenstein, M., 2014. Age-related effects on ERP and oscillatory EEG-dynamics in a 2-back task. J. Psychophysiol. 10.1027/0269-8803/a000123

Haaland, K.Y., Vranes, L.F., Goodwin, J.S., Garry, P.J., 1987. Wisconsin Card Sort Test Performance in a Healthy Elderly Population. J. Gerontol. 42, 345–346. 10.1093/geronj/42.3.345

Hakun, J.G., Zhu, Z., Brown, C.A., Johnson, N.F., Gold, B.T., 2015. Longitudinal alterations to brain function, structure, and cognitive performance in healthy older adults: A fMRI-DTI study. Neuropsychologia 71, 225–235. 10.1016/j.neuropsychologia.2015.04.008

He, M., Liu, F., Nummenmaa, A., Hämäläinen, M., Dickerson, B.C., Purdon, P.L., 2021. Age-Related EEG Power Reductions Cannot Be Explained by Changes of the Conductivity Distribution in the Head Due to Brain Atrophy. Front. Aging Neurosci. 13, 632310. 10.3389/fnagi.2021.632310

Huizeling, E., Wang, H., Holland, C., Kessler, K., 2021. Changes in theta and alpha oscillatory signatures of attentional control in older and middle age. Eur. J. Neurosci. 54, 4314–4337. 10.1111/ejn.15259

JASP Team, 2024. JASP.

Jernigan, T.L., Archibald, S.L., Fennema-Notestine, C., Gamst, A.C., Stout, J.C., Bonner, J., Hesselink, J.R., 2001. Effects of age on tissues and regions of the cerebrum and cerebellum. Neurobiol. Aging 22, 581–594. 10.1016/S0197-4580(01)00217-2

Kaiser, J., Simon, N.A., Sauseng, P., Schütz-Bosbach, S., 2019. Midfrontal neural dynamics distinguish between general control and inhibition-specific processes in the stopping of motor actions. Sci. Rep. 9, 13054. 10.1038/s41598-019-49476-4

Karayanidis, F., Whitson, L.R., Heathcote, A., Michie, P.T., 2011. Variability in Proactive and Reactive Cognitive Control Processes Across the Adult Lifespan. Front. Psychol. 2. 10.3389/fpsyg.2011.00318

Karbach, J., Kray, J., 2009. How useful is executive control training? Age differences in near and far transfer of task-switching training. Dev. Sci. 12, 978–990. 10.1111/j.1467-7687.2009.00846.x

Kennedy, K.M., Raz, N., 2009. Aging white matter and cognition: Differential effects of regional variations in diffusion properties on memory, executive functions, and speed. Neuropsychologia 47, 916–927. 10.1016/j.neuropsychologia.2009.01.001

Koen, J.D., Rugg, M.D., 2019. Neural Dedifferentiation in the Aging Brain. Trends Cogn. Sci. 23, 547–559. 10.1016/j.tics.2019.04.012

Kolev, V., Falkenstein, M., Yordanova, J., 2024. A distributed theta network of error generation and processing in aging. Cogn. Neurodyn. 18, 447–459. 10.1007/s11571-023-10018-4

Kolev, V., Falkenstein, M., Yordanova, J., 2005. Aging and Error Processing: Time-Frequency Analysis of Error-Related Potentials. J. Psychophysiol. 19, 289–297. https://psycnet.apa.org/doi/10.1027/0269-8803.19.4.289

Kray, J., 2006. Task-set switching under cue-based versus memory-based switching conditions in younger and older adults. Brain Res. 1105, 83–92. 10.1016/j.brainres.2005.11.016

Kray, J., Li, K.Z.H., Lindenberger, U., 2002. Age-Related Changes in Task-Switching Components: The Role of Task Uncertainty. Brain Cogn. 49, 363–381. 10.1006/brcg.2001.1505

Kray, J., Lindenberger, U., 2000. Adult age differences in task switching. Psychol. Aging 15, 126–147. 10.1037/0882-7974.15.1.126

Küçük, K.M., Mathes, B., Schmiedt-Fehr, C., Başar-Eroğlu, C., 2023. Aging attenuated theta response during multistable perception. Psychophysiology 60, e14286. 10.1111/psyp.14286

Lehrl, S., Triebig, G., Fischer, B., 1995. Multiple choice vocabulary test MWT as a valid and short test to estimate premorbid intelligence. Acta Neurol. Scand. 91, 335–345. 10.1111/j.1600-0404.1995.tb07018.x

Liesefeld, H.R., Janczyk, M., 2019. Combining speed and accuracy to control for speed-accuracy trade-offs(?). Behav. Res. Methods 51, 40–60. 10.3758/s13428-018-1076-x

Luck, S.J., 2014. An introduction to the event-related potential technique, Second edition. ed. The MIT Press, Cambridge.

McDermott, T.J., Wiesman, A.I., Proskovec, A.L., Heinrichs-Graham, E., Wilson, T.W., 2017. Spatiotemporal oscillatory dynamics of visual selective attention during a flanker task. NeuroImage 156, 277–285. 10.1016/j.neuroimage.2017.05.014

Meiran, N., Gotler, A., Perlman, A., 2001. Old Age Is Associated With a Pattern of Relatively Intact and Relatively Impaired Task-Set Switching Abilities. J. Gerontol. Ser. B 56, P88–P102. 10.1093/geronb/56.2.P88

Monsell, S., 2003. Task switching. Trends Cogn. Sci. 7, 134–140. 10.1016/S1364-6613(03)00028-7

Morales, S., Bowers, M.E., 2022. Time-frequency analysis methods and their application in developmental EEG data. Dev. Cogn. Neurosci. 54, 101067. 10.1016/j.dcn.2022.101067

Moretti, D.V., Miniussi, C., Frisoni, G.B., Geroldi, C., Zanetti, O., Binetti, G., Rossini, P.M., 2007. Hippocampal atrophy and EEG markers in subjects with mild cognitive impairment. Clin. Neurophysiol. 118, 2716–2729. 10.1016/j.clinph.2007.09.059

Nigbur, R., Ivanova, G., Stürmer, B., 2011. Theta power as a marker for cognitive interference. Clin. Neurophysiol. 122, 2185–2194. 10.1016/j.clinph.2011.03.030

Oh, A., Vidal, J., Taylor, M.J., Pang, E.W., 2014. Neuromagnetic correlates of intra- and extra-dimensional set-shifting. Brain Cogn. 86, 90–97. 10.1016/j.bandc.2014.02.006

Oostenveld, R., Fries, P., Maris, E., Schoffelen, J.-M., 2011. FieldTrip: Open Source Software for Advanced Analysis of MEG, EEG, and Invasive Electrophysiological Data. Comput. Intell. Neurosci. 2011, 1–9. 10.1155/2011/156869

Owen, A.M., Roberts, A.C., Polkey, C.E., Sahakian, B.J., Robbins, T.W., 1991. Extra-dimensional versus intra-dimensional set shifting performance following frontal lobe excisions, temporal lobe excisions or amygdalo-hippocampectomy in man. Neuropsychologia 29, 993–1006. 10.1016/0028-3932(91)90063-E

Panitz, C., Keil, A., Mueller, E.M., 2019. Extinction-resistant attention to long-term conditioned threat is indexed by selective visuocortical alpha suppression in humans. Sci. Rep. 9, 15809. 10.1038/s41598-019-52315-1

Pardo, J.V., Lee, J.T., Sheikh, S.A., Surerus-Johnson, C., Shah, H., Munch, K.R., Carlis, J.V., Lewis, S.M., Kuskowski, M.A., Dysken, M.W., 2007. Where the brain grows old: decline in anterior cingulate and medial prefrontal function with normal aging. NeuroImage 35, 1231–7. 10.1016/j.neuroimage.2006.12.044

R Core Team, 2022.R: A Language and Environment for Statistical Computing. R Studio Team, 2020. RStudio: Integrated Development Environment for R.

Ratcliffe, O., Shapiro, K., Staresina, B.P., 2022. Fronto-medial theta coordinates posterior maintenance of working memory content. Curr. Biol. 32, 2121–2129.e3. 10.1016/j.cub.2022.03.045

Reuter-Lorenz, P.A., 2002. New visions of the aging mind and brain. Trends Cogn. Sci. 6, 394–400. 10.1016/S1364-6613(02)01957-5

Reynolds, C.R., James, B.J., Mayfield, J., 1995. Baseline Performance of Normal Elderly on a Verbal Measurement of Set-Shifting and Executive Function. Arch. Clin. Neuropsychol. 4, 383. 10.1093/arclin/10.4.383

Richter, A., Soch, J., Kizilirmak, J.M., Fischer, L., Schütze, H., Assmann, A., Behnisch, G., Feldhoff, H., Knopf, L., Raschick, M., Schult, A., Seidenbecher, C.I., Yakupov, R., Düzel, E., Schott, B.H., 2023. Single-value scores of memory-related brain activity reflect dissociable neuropsychological and anatomical signatures of neurocognitive aging. Hum. Brain Mapp. 44, 3283–3301. 10.1002/hbm.26281

Ridderinkhof, K.R., Span, M.M., van der Molen, M.W., 2002. Perseverative Behavior and Adaptive Control in Older Adults: Performance Monitoring, Rule Induction, and Set Shifting. Brain Cogn. 49, 382–401. 10.1006/brcg.2001.1506

Sahakian, B.J., Owen, A.M., 1992. Computerized assessment in neuropsychiatry using CANTAB: discussion paper. J. R. Soc. Med. 85, 399–402.

Sauseng, P., Klimesch, W., Freunberger, R., Pecherstorfer, T., Hanslmayr, S., Doppelmayr, M., 2006. Relevance of EEG alpha and theta oscillations during task switching. Exp. Brain Res. 170, 295–301. 10.1007/s00221-005-0211-y

Schott, B.H., Niklas, C., Kaufmann, J., Bodammer, N.C., Machts, J., Schutze, H., Duzel, E., 2011. Fiber density between rhinal cortex and activated ventrolateral prefrontal regions predicts episodic memory performance in humans. Proc. Natl. Acad. Sci. 108, 5408–5413. 10.1073/pnas.1013287108

Schwarzkopf, D., de Haas, B., Rees, G., 2012. Better Ways to Improve Standards in Brain-Behavior Correlation Analysis. Front. Hum. Neurosci. 6. 10.3389/fnhum.2012.00200

Seeley, W.W., 2019. The Salience Network: A Neural System for Perceiving and Responding to Homeostatic Demands. J. Neurosci. 39, 9878. 10.1523/JNEUROSCI.1138-17.2019

Starns, J.J., Ratcliff, R., 2010. The effects of aging on the speed–accuracy compromise: Boundary optimality in the diffusion model. Psychol. Aging 25, 377–390. 10.1037/a0018022

Steyvers, M., Hawkins, G.E., Karayanidis, F., Brown, S.D., 2019. A large-scale analysis of task switching practice effects across the lifespan. Proc Natl Acad Sci U A 116, 17735–17740. 10.1073/pnas.1906788116

Stolz, C., Pickering, A.D., Mueller, E.M., 2023. Dissociable feedback valence effects on frontal midline theta during reward gain versus threat avoidance learning. Psychophysiology 60. 10.1111/psyp.14235

Strobach, T., Liepelt, R., Schubert, T., Kiesel, A., 2012. Task switching: effects of practice on switch and mixing costs. Psychol. Res. 76, 74–83. 10.1007/s00426-011-0323-x

The MathWorks Inc., ;, 2021. MATLAB Version 9.11.0.1809720 (R2021b).

Tsujimoto, T., Shimazu, H., Isomura, Y., 2006. Direct Recording of Theta Oscillations in Primate Prefrontal and Anterior Cingulate Cortices. J. Neurophysiol. 95, 2987–3000. 10.1152/jn.00730.2005

van de Vijver, I., Cohen, M.X., Ridderinkhof, K.R., 2014. Aging affects medial but not anterior frontal learning-related theta oscillations. Neurobiol. Aging 35, 692–704. 10.1016/j.neurobiolaging.2013.09.006

van de Vijver, I., Ridderinkhof, K.R., Cohen, M.X., 2011. Frontal Oscillatory Dynamics Predict Feedback Learning and Action Adjustment. J. Cogn. Neurosci. 23, 4106–4121. 10.1162/jocn_a_00110

Velanova, K., Lustig, C., Jacoby, L.L., Buckner, R.L., 2006. Evidence for Frontally Mediated Controlled Processing Differences in Older Adults. Cereb. Cortex 17, 1033–1046. 10.1093/cercor/bhl013

Watson, T.D., Azizian, A., Squires, N.K., 2006. Event-related potential correlates of extradimensional and intradimensional set-shifts in a modified Wisconsin Card Sorting Test. Brain Res. 1092, 138–151. 10.1016/j.brainres.2006.03.098

Wecker, N.S., Kramer, J.H., Hallam, B.J., Delis, D.C., 2005. Mental Flexibility: Age Effects on Switching. Neuropsychology 19, 345–352. 10.1037/0894-4105.19.3.345

Whitson, L.R., Karayanidis, F., Fulham, R., Provost, A., Michie, P.T., Heathcote, A., Hsieh, S., 2014. Reactive control processes contributing to residual switch cost and mixing cost across the adult lifespan. Front. Psychol. 5. 10.3389/fpsyg.2014.00383

Winkler, I., Debener, S., Müller, K.-R., Tangermann, M., 2015. On the influence of high-pass filtering on ICA-based artifact reduction in EEG-ERP. Presented at the 2015 37th Annual International Conference of the IEEE Engineering in Medicine and Biology Society (EMBC), pp. 4101– 4105. 10.1109/EMBC.2015.7319296

Wylie, G., Allport, A., 2000. Task switching and the measurement of “switch costs.” Psychol. Res. 63, 212–233. 10.1007/s004269900003

Yener, G., Hünerli-Gündüz, D., Yıldırım, E., Aktürk, T., Başar-Eroğlu, C., Bonanni, L., Del Percio, C., Farina, F., Ferri, R., Güntekin, B., Hajós, M., Ibáñez, A., Jiang, Y., Lizio, R., Lopez, S., Noce, G., Parra, M.A., Randall, F., Stocchi, F., Babiloni, C., 2022. Treatment effects on event-related EEG potentials and oscillations in Alzheimer’s disease. Int. J. Psychophysiol. 177, 179–201. 10.1016/j.ijpsycho.2022.05.008

Yeung, M.K., Han, Y.M.Y., Sze, S.L., Chan, A.S., 2016. Abnormal frontal theta oscillations underlie the cognitive flexibility deficits in children with high-functioning autism spectrum disorders. Neuropsychology 30, 281–295. 10.1037/neu0000231

Yordanova, J., Falkenstein, M., Kolev, V., 2020. Aging-related changes in motor response-related theta activity. Int. J. Psychophysiol. 153, 95–106. 10.1016/j.ijpsycho.2020.03.005

Zelazo, P.D., Craik, F.I.M., Booth, L., 2004. Executive function across the life span. Acta Psychol. (Amst.) 115, 167–183. 10.1016/j.actpsy.2003.12.005

Ziegler, D.A., Piguet, O., Salat, D.H., Prince, K., Connally, E., Corkin, S., 2010. Cognition in healthy aging is related to regional white matter integrity, but not cortical thickness. Neurobiol Aging 31, 1912–26. 10.1016/j.neurobiolaging.2008.10.015

