## Supplementary Material for "Frontal Theta Oscillations and Cognitive Flexibility: Age-Related Modulations in EEG Activity"

### S1. Supplementary Tables

**Supplementary Table 1:** Subject-specific criteria for raw data inspection and data rejection.

| Subject Nr. | Gradient_max | Channel with highest rejection | Percentage rejected from continuous data |
|---|---|---|---|
| 1 | 70 | FT8 | 16.0 % |
| 2 | 60 | FC5 | 8.3 % |
| 3 | 50 | FC4 | 2.9 % |
| 4 | 60 | Oz | 2.0 % |
| 5 | 40 | FC6 | 6.3 % |
| 6 | 50 | C5 | 9.9 % |
| 7 | 40 | T7 | 0.5 % |
| 8 | 50 | POz | 6.1 % |
| 9 | 40 | FC5 | 7.0 % |
| 10 | 40 | FC5 | 2.3 % |
| 11 | 40 | T7 | 4.4 % |
| 12 | 50 | O1 | 1.0 % |
| 13 | 50 | F4 | 9.9 % |
| 14 | 50 | FC4 | 13.5 % |
| 15 | 60 | T7 | 6.9 % |
| 16 | 50 | Oz | 1.9 % |
| 17 | 40 | TP8 | 7.9 % |
| 18 | 60 | FP2 | 4.5 % |
| 19 | 40 | T7 | 4.5 % |
| 20 | 40 | PO9 | 7.9 % |
| 21 | 50 | TP7 | 6.0 % |
| 22 | 70 | T7 | 2.3 % |
| 23 | 40 | T7 | 6.5 % |
| 24 | 50 | FC5 | 6.4 % |
| 25 | 60 | FT7 | 10 % |
| 26 | 50 | CP3 | 2.7 % |
| 27 | 40 | AF7 | 7.1 % |
| 28 | 70 | FC6 | 5.4 % |
| 29 | 40 | T7 | 10 % |
| 30 | 50 | TP8 | 11.1 % |
| 31 | 50 | F8 | 0.4 % |
| 32 | 40 | T7 | 3.3 % |
| 33 | 40 | FC5 | 4.8 % |
| 34 | 40 | Fp1 | 6.5 % |
| 35 | 40 | Fp2 | 9.4 % |
| 36 | 60 | FC6 | 3.3 % |
| 37 | 50 | AF7 | 8.8 % |
| 38 | 60 | P7 | 5.8 % |
| 39 | 40 | C6 | 14.4 % |

**Gradient_max:** maximum allowed voltage step on the gradient. Gradient before: 400 ms, Gradient after: 400 ms.

**Supplementary Table 2:** Name of channels that were interpolated at least once followed by the number of young and older subjects for which the interpolation was performed.

| Channel | No. of young subjects | No. of older subjects |
|---|---|---|
| TP9 | 1 | 1 |
| TP8 | 1 | - |
| PO9 | 1 | - |

### S2. IDED Supplementary Figure

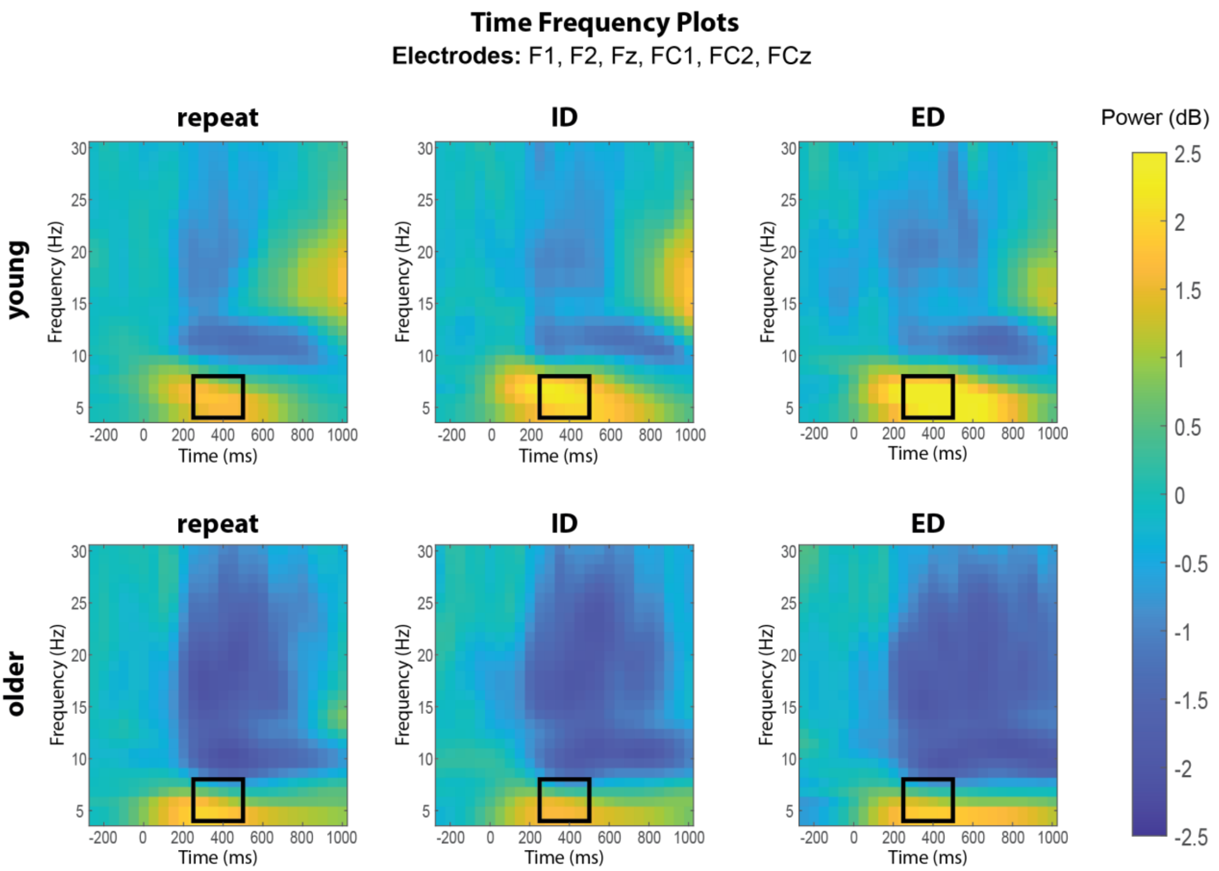

**Supplementary Figure 1: Time Frequency Plots**. Power of respective frequencies over time averaged over F1, F2, Fz, FC1, FC2, FCz. The chosen frequency range (4-8Hz) and time range (250 to 500 ms) are included in the black outlined square.

### S3. Older participants have difficulties in attentional set-shifting

##### S3.1Attentional Set Shifting Task (ASST)

The Attentional Set Shifting Task (ASST) is a paradigm in which participants are asked to discern, which stimulus is correct. The correct stimulus of each stage is predetermined and the participants have to discover it by trial and error. In each trial, a fixation cross is first presented for a pseudorandomized interval of 800 to 1300 ms. Next, two stimuli appear, one to the left and one to the right side of the fixation cross. The participant is instructed to choose the left or right stimulus by pressing the respective mouse button. After button press or a response time cut-off of 3000 ms, a black screen is shown for 800 to 1000 ms, followed by presentation of the feedback (a green circle for correct responses or a red circle for incorrect responses) for 1000 ms (**Supplementary Figure 2**A).

The ASST consists of multiple stages, each composed of several trials in which the correct stimulus remains unchanged. Eight consecutive correct responses were defined as criterion that the correct stimulus was recognized and lead to the advancement to the following stage, at which the correct stimulus changes and has to be discerned once more. The task is terminated when a participant successfully completes all stages or when 50 trials in one stage were reached without successfully recognizing the correct stimulus. Notably, the participants are beforehand neither informed about the number of correct trials required to move to the next stage, nor about the total number of stages or the relevant dimensions.

With the exception of the first stage, the participants are presented with stimuli composed of two superimposed elements, a grey shape and white line pattern, resulting in a compound stimulus, with a shape and a line dimension. Within each stage, one exemplar of one dimension is the correct one whereas the other dimension is irrelevant. Additionally, in the trials within a stage the same exemplars are used but their configuration and position change independently from trial to trial (for more information see: Dickson et al. 2014).

The ASST task consists of eight stages with increasing shift difficulty (Supplementary Figure 2B). In the first stage (simple discrimination, SD), simple stimuli are presented of either the line or shape dimension. This is followed by the compound discrimination (CD) stage, in which the compound stimuli first appear. Subsequently the set-shifting stages are: compound discrimination reversal (CDR), intra-dimensional shift 1 (IDS1), intra-dimensional shift 1 reversal (IDS1R), intra-dimensional shift 2 (IDS2), intra-dimensional shift 2 reversal (IDS2R) and extra-dimensional shift (EDS). During reversal stages (CDR, IDS1R, IDS2R) the previously incorrect exemplar of the relevant dimension becomes correct. In the IDS1 and IDS2 stages, new stimuli are presented but the correct exemplar belongs in the same dimension as in the previous stage. Lastly, in the extra-dimensional shift stage, the correct exemplar belongs in the previously irrelevant dimension.

**S3.2 Statistical Analysis of the ASST**

Behavioural performance in the ASST was evaluated by estimating a survival curve using the Kaplan-Meier method. Each stage of the paradigm was evaluated as an event, and the probability of “survival” was defined as the probability of successfully completing all stages of the paradigm. The estimated probability of survival is reported together with the standard error.

**S3.2 Results from the ASST**

Among 20 young participants, only one failed to complete the ASST, whereas 14 of 19 older participants did not finish the task. The majority of them failed in the ED shift stage (n=9). A survival analysis revealed significant differences in task completion between the two groups ($X^2(1, N = 39) = 17.20$, $p < .001$) with young participants completing the task at an estimated probability of 0.95 ± 0.05 (Standard Error) at the last stage (Supplementary Figure 3).

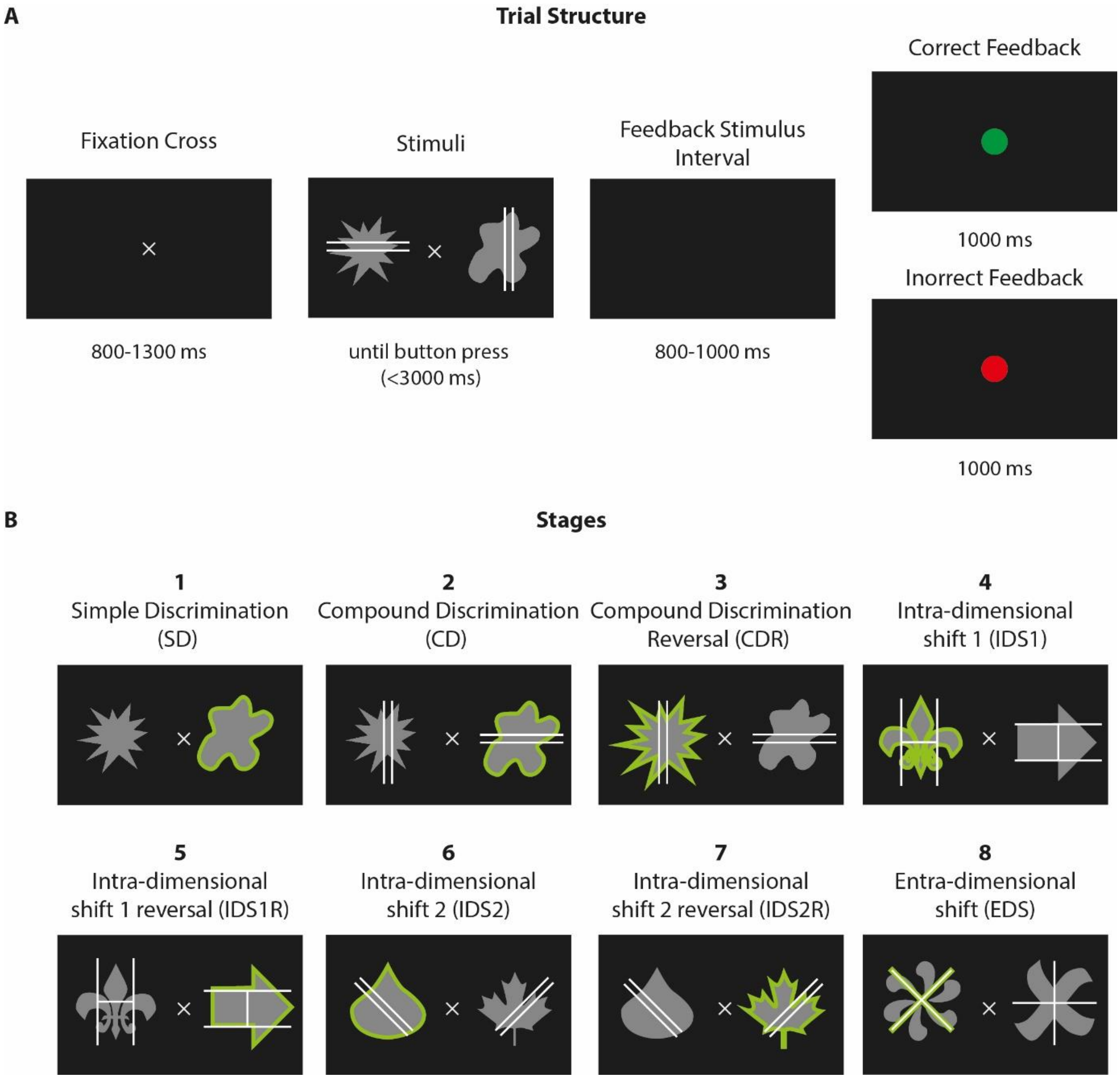

**Supplementary Figure 2: A:** Trial Structure of the ASST. After the presentation of the fixation cross for a pseudorandomized interval of 800 to 1300 ms the two stimuli appear. The participant clicks the left or right mouse button to select the left or right stimulus, respectively. A black screen is visible for 800 to 1000 ms after the button is pressed or after 3000 ms without response. This is followed by the feedback for 1000 ms: green circle for correct choice and red circle for incorrect; **B:** Exemplary visualization of the 8 ASST stages. The green shade indicates the pattern that elicits the correct response in each stage (not visible to participants). The participants are required to discern the correct stimulus through trial and error. A new stage automatically starts after 8 consecutive correct responses. In each stage only one pair of compound stimuli is presented (this equates to two exemplars of each dimension). However, the stimulus position and configuration are independently randomized in each trial. For more information see Dickson et al. (2014).

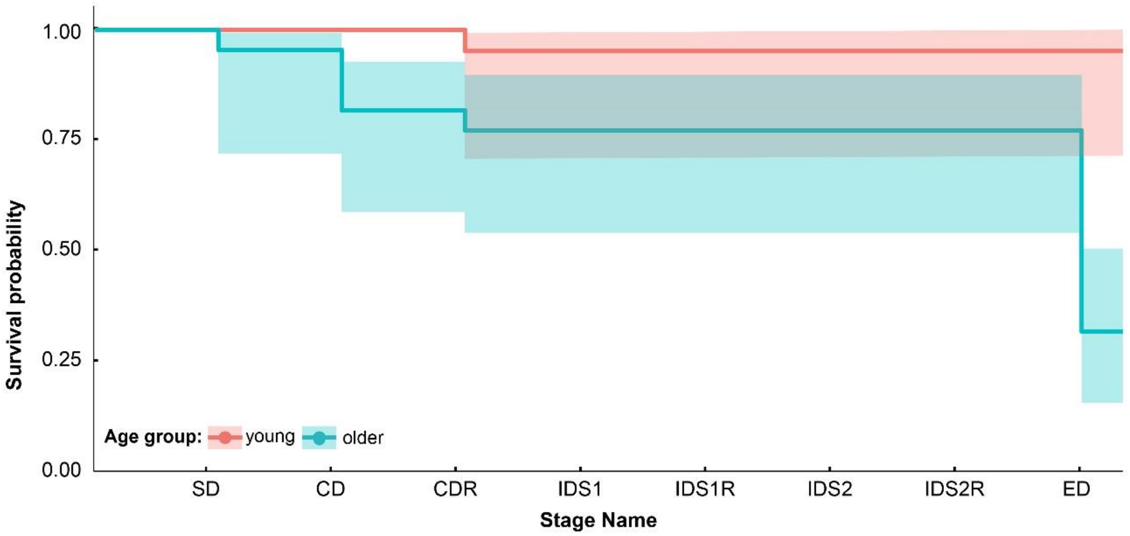

**Supplementary Figure 3:** Probability to complete the Attentional Set Shifting Task (ASST) for the two age groups, plotted as survival curve. There is a significant difference in task completion rates between the two age groups (X2(1, N = 39) = 17.20, p < .001). The majority of older individuals fail to complete the ASST at the ED stage. The shaded areas represent Standard Errors of the survival curves. SD: Simple Discrimination; CD: Compound Discrimination; CDR: Compound Discrimination Reversal; IDS1: Intra-dimensional shift 1; IDS1R: Intra-dimensional shift 1 reversal; IDS2: Intra-dimensional shift 2; IDS2R: Intra-dimensional shift 2 reversal; EDS: Extra-dimensional.

#### S3.3 Discussion of the ASST results

As evident from the results, the majority of older adults failed to successfully solve the ED stage. This finding is in line with previous studies (De Luca et al., 2003; Owen et al., 1991; Zelazo et al., 2004) and indicates that older adults have more difficulty solving the ED stage in this particular task. In contrast to the IDED, the correct answer of each stage of the ASST is not apparent to the participant, but has to be discerned via trial and error. No external input is given from the experimenter in regards to the relevant stimulus dimensions and strategies that may be used. It therefore appears here that older individuals fail to inhibit the original set associations and switch to a new dimension n the ED shift. On the other hand, most young participants recognize that possibility and successfully complete the paradigm. This difference between age groups may count as an indicator that many older adults have difficulties in ED shifting.

As of note, five of the older adults failed the task in the early stages. This early failure may be unrelated to cognitive flexibility and may reflect unrelated cognitive processes in which these particular individuals may have deficits in. Based on our criteria for cognitive screening, all participants, however, reached the cognitive criterium of an MMSE score above 24 (Creavin et al., 2016), with an average test-score of 28.2 ± 1.3. The participant with the lowest MMSE score (26) additionally took part in a more extensive cognitive evaluation of working memory, alertness and executive function (Test Battery for Attention (TAP); Zimmermann & Fimm, 1992; Verbal Learning and Memory Test (VLMT); Helmstaedter et al., 2001; Logical Memory subtest from the Wechsler Memory Scale (WMS); Härting et al., 2000; Flanker task, N-back task). The results in those tasks did not differ from the average ± 3 standard deviation of participants of similar to other participants of the same age group. in our previously established cohort (Richter et al., 2023).

Notably, there is a striking performance difference between the ASST and the IDED in the older participants of our study, as many participants failed the ED stage of the ASST, yet all older participants showcased low error rates. We explain this through the substantial difference in task difficulty. Given its structure, the IDED is a very easy task and the answer is always apparent. In contrast to that, the ASST entails more trial-and-error and shifts to dimensions that may not be apparent to the participants. Additionally, in contrast to the IDED, the participants did not receive any hints on which the possible dimensions were. Thus, even participants who failed to complete the ASST were still able to finish the IDED.

### S4. Robust ANOVA to control for non-normality on reaction times and error rates of the IDED task

The mixed robust ANOVAs on RTs and error rates of the intra-dimensional/extra-dimensional (IDED) task were performed as described in Mair and Wilcox (2020) using the WRS2 package. The robust ANOVAs were performed based on 20% trimmed means (function: *rmanova*). In case of significant effects of condition, the function *rmmcp* for corresponding *post-hoc* tests were used. In case of significant effects of condition, *post-hoc* tests were not necessary, since only two factors were evaluated (young vs older). In both situations, effect size was calculated as robust standardized difference similar to Cohen's d (Algina et al., 2005) for dependent variables (function: *dep.effect*) and independent variables (function: *akp.effect*) respectively.

When comparing error rates, we found a significant effect of condition ($F$ (2, 18.54) = 9.889, $p$ = .001) and neither an effect of age group ($F$ (1, 21.32) = 0.445, $p$ = .512) nor an interaction between the two factors was significant ($F$ (2, 18.54) = 0.016, $p$ = .984). *Post-hoc* tests within condition revealed significant differences in error rates amongst all conditions (ED vs ID: $\hat{\psi}$ = 0.013, $p$ = .005, *AKP* = 0.530; ED vs repeat: $\hat{\psi}$ = 0.021, $p$ < .001, *AKP* = 0.841; ID vs repeat: $\hat{\psi}$ = 0.006, $p$ = .026, *AKP* = 0.405).

### S5. Multiple Comparisons after three-way ANOVA

| Age group | Electrode | Contrast | *b* | *SE* | *t* | $P_{adj}$ | sig |
|---|---|---|---|---|---|---|---|
| young | Fz | repeat - ID | -0.57 | 0.14 | -4.00 | < **.001** | *** |
| | | repeat - ED | -0.82 | 0.13 | -6.49 | < **.001** | *** |
| | | ID - ED | -0.25 | 0.14 | -1.84 | .073 | |
| | FCz | repeat - ID | -0.63 | 0.15 | -4.32 | < **.001** | *** |
| | | repeat - ED | -1.07 | 0.12 | -8.70 | < **.001** | *** |
| | | ID - ED | -0.44 | 0.13 | -3.37 | **.002** | ** |
| | Cz | repeat - ID | -0.55 | 0.15 | -3.68 | **.001** | ** |
| | | repeat - ED | -0.98 | 0.13 | -7.75 | < **.001** | *** |
| | | ID - ED | -0.42 | 0.14 | -3.01 | **.005** | ** |
| | CPz | repeat - ID | -0.45 | 0.15 | -3.00 | **.007** | ** |
| | | repeat - ED | -0.67 | 0.13 | -5.05 | < **.001** | *** |
| | | ID - ED | -0.22 | 0.15 | -1.48 | .148 | |
| | Pz | repeat - ID | -0.34 | 0.17 | -2.03 | .074 | |
| | | repeat - ED | -0.38 | 0.14 | -2.69 | **.032** | * |
| | | ID - ED | -0.05 | 0.16 | -0.30 | .770 | |
| | POz | repeat - ID | -0.32 | 0.17 | -1.95 | .178 | |
| | | repeat - ED | -0.24 | 0.16 | -1.49 | .216 | |
| | | ID - ED | 0.08 | 0.17 | 0.47 | .644 | |
| | Oz | repeat - ID | -0.28 | 0.16 | -1.81 | .233 | |
| | | repeat - ED | -0.12 | 0.14 | -0.87 | .233 | |
| | | ID - ED | 0.16 | 0.17 | 0.97 | .392 | |
| older | Fz | repeat - ID | 0.07 | 0.15 | 0.49 | .919 | |
| | | repeat - ED | 0.08 | 0.13 | 0.66 | .919 | |
| | | ID - ED | 0.01 | 0.14 | 0.10 | .919 | |
| | FCz | repeat - ID | 0.03 | 0.15 | 0.21 | .832 | |
| | | repeat - ED | 0.10 | 0.13 | 0.80 | .832 | |
| | | ID - ED | 0.07 | 0.13 | 0.51 | .832 | |
| | Cz | repeat - ID | 0.03 | 0.16 | 0.21 | .832 | |
| | | repeat - ED | 0.10 | 0.13 | 0.81 | .832 | |
| | | ID - ED | 0.07 | 0.14 | 0.50 | .832 | |
| | CPz | repeat - ID | 0.05 | 0.16 | 0.32 | .936 | |
| | | repeat - ED | -0.01 | 0.14 | -0.08 | .936 | |
| | | ID - ED | -0.06 | 0.15 | -0.41 | .936 | |
| | Pz | repeat - ID | -.01 | 0.17 | 0.07 | .943 | |
| | | repeat - ED | -0.09 | 0.15 | -0.60 | .827 | |
| | | ID - ED | -0.10 | 0.17 | -0.60 | .827 | |
| | POz | repeat - ID | -0.01 | 0.17 | -0.07 | .945 | |
| | | repeat - ED | -0.17 | 0.17 | -1.00 | .591 | |
| | | ID - ED | -0.15 | 0.18 | -0.86 | .591 | |
| | Oz | repeat - ID | -0.15 | 0.16 | -0.94 | .370 | |
| | | repeat - ED | -0.31 | 0.14 | -2.12 | .122 | |
| | | ID - ED | -0.16 | 0.17 | -0.91 | .370 | |

***SE*:** Standard error, ***$P_{adj}$*:** FDR corrected p value, ***sig***: * for $p \leq .05$ , ** for $p \leq .005$ and *** for $p \leq .001$.

### S6. Supplementary References

Algina, J., Keselman, H.J., Penfield, R.D., 2005. An alternative to Cohen's standardized mean difference effect size: a robust parameter and confidence interval in the two independent groups case. Psychol Methods 10(3), 317-328. https://doi.org/10.1037/1082-989x.10.3.317.

Alperin, B.R., Haring, A.E., Zhuravleva, T.Y., Holcomb, P.J., Rentz, D.M., Daffner, K.R., 2013. The Dissociation between Early and Late Selection in Older Adults. Journal of Cognitive Neuroscience 25(12), 2189-2206. https://doi.org/10.1162/jocn_a_00456.

Barceló, F., 2003. The Madrid card sorting test (MCST): a task switching paradigm to study executive attention with event-related potentials. Brain Research Protocols 11(1), 27–37. https://doi.org/10.1016/S1385-299X(03)00013-8.

Creavin, S.T., Wisniewski, S., Noel-Storr, A.H., Trevelyan, C.M., Hampton, T., Rayment, D., Thom, V.M., Nash, K.J.E., Elhamoui, H., Milligan, R., et al., 2016. Mini-Mental State Examination (MMSE) for the detection of dementia in clinically unevaluated people aged 65 and over in community and primary care populations. Cochrane Database of Systematic Reviews(1). https://doi.org/10.1002/14651858.cd011145.pub2.

de Jong, H.L., Kok, A., van Rooy, J.C.G.M., 1988. Early and Late Selection in Young and Old Adults: An Event-Related Potential Study. Psychophysiology 25(6), 657-671. https://doi.org/10.1111/j.1469-8986.1988.tb01904.x.

De Luca, C.R., Wood, S.J., Anderson, V., Buchanan, J.-A., Proffitt, T.M., Mahony, K., Pantelis, C., 2003. Normative Data From the Cantab. I: Development of Executive Function Over the Lifespan. Journal of Clinical and Experimental Neuropsychology 25(2), 242-254. https://doi.org/10.1076/jcen.25.2.242.13639.

Dickson, P.E., Calton, M.A., Mittleman, G., 2014. Performance of C57BL/6J and DBA/2J mice on a touchscreen-based attentional set-shifting task. Behavioural brain research 261, 158–170. https://doi.org/10.1016/j.bbr.2013.12.015.

Duncan, C.C., Barry, R.J., Connolly, J.F., Fischer, C., Michie, P.T., Näätänen, R., Polich, J., Reinvang, I., Van Petten, C., 2009. Event-related potentials in clinical research: Guidelines for eliciting, recording, and quantifying mismatch negativity, P300, and N400. Clinical Neurophysiology 120(11), 1883-1908. https://doi.org/10.1016/j.clinph.2009.07.045.

Eppinger, B., Kray, J., Mecklinger, A., John, O., 2007. Age differences in task switching and response monitoring: Evidence from ERPs. Biological Psychology 75(1), 52-67. https://doi.org/10.1016/j.biopsycho.2006.12.001.

Fernandes, L.S., Ferreira, D.S., Almeida, P.R., Dias, N.S., 2015. Aging and attentional set shifting on WCST: An event-related EEG study, 2015 7th International IEEE/EMBS Conference on Neural Engineering (NER). pp. 1088-1091. https://doi.org/10.1109/NER.2015.7146817.

Gajewski, P.D., Falkenstein, M., 2014. Age-related effects on ERP and oscillatory EEG-dynamics in a 2-back task. Journal of Psychophysiology. https://doi.org/10.1027/0269-8803/a000123.

Härting, C., Markowitsch, H., Neufeld, H., Calabrese, P., Deisinger, K., Kessler, J., 2000. WMS-R–Manual. Bern: Hans Huber Verlag.

Helmstaedter, C., Lendt, M., Lux, S., 2001. Verbaler Lern-und Merkfähigkeitstest: VLMT; Manual. Beltz-test.

Hillyard, S.A., Anllo-Vento, L., 1998. Event-related brain potentials in the study of visual selective attention. Proceedings of the National Academy of Sciences 95(3), 781-787. https://doi.org/10.1073/pnas.95.3.781.

Hirayasu, Y., Samura, M., Ohta, H., Ogura, C., 2000. Sex effects on rate of change of P300 latency with age. Clinical Neurophysiology 111(2), 187-194. https://doi.org/10.1016/S1388-2457(99)00233-3.

Jost, K., Mayr, U., Rösler, F., 2008. Is task switching nothing but cue priming? Evidence from ERPs. Cognitive, Affective, & Behavioral Neuroscience 8(1), 74-84. https://doi.org/10.3758/CABN.8.1.74.

Karayanidis, F., Coltheart, M., Michie, P.T., Murphy, K., 2003. Electrophysiological correlates of anticipatory and poststimulus components of task switching. Psychophysiology 40(3), 329-348. https://doi.org/10.1111/1469-8986.00037.

Karayanidis, F., Jamadar, S., Ruge, H., Phillips, N., Heathcote, A., Forstmann, B.U., 2010. Advance preparation in task-switching: converging evidence from behavioral, brain activation, and model-based approaches. Frontiers in Psychology 1, 25. https://doi.org/10.3389/fpsyg.2010.00025.

Karayanidis, F., Mansfield, E.L., Galloway, K.L., Smith, J.L., Provost, A., Heathcote, A., 2009. Anticipatory reconfiguration elicited by fully and partially informative cues that validly predict a switch in task. Cogn Affect Behav Neurosci 9(2), 202-215. https://doi.org/10.3758/CABN.9.2.202.

Karayanidis, F., Whitson, L.R., Heathcote, A., Michie, P.T., 2011. Variability in Proactive and Reactive Cognitive Control Processes Across the Adult Lifespan. Frontiers in Psychology 2. https://doi.org/10.3389/fpsyg.2011.00318.

Keil, A., Müller, M.M., 2010. Feature selection in the human brain: Electrophysiological correlates of sensory enhancement and feature integration. Brain Research 1313, 172-184. https://doi.org/10.1016/j.brainres.2009.12.006.

Kopp, B., Lange, F., 2013. Electrophysiological indicators of surprise and entropy in dynamic task-switching environments. Frontiers in Human Neuroscience 7(300). https://doi.org/10.3389/fnhum.2013.00300.

Lange, F., Seer, C., Müller, D., Kopp, B., 2015. Cognitive caching promotes flexibility in task switching: evidence from event-related potentials. Scientific reports 5, 17502. https://doi.org/10.1038/srep17502.

Mair, P., Wilcox, R., 2020. Robust statistical methods in R using the WRS2 package. Behavior Research Methods 52(2), 464-488. https://doi.org/10.3758/s13428-019-01246-w.

Nagahama, Y., Okada, T., Katsumi, Y., Hayashi, T., Yamauchi, H., Oyanagi, C., Konishi, J., Fukuyama, H., Shibasaki, H., 2001. Dissociable mechanisms of attentional control within the human prefrontal cortex. Cerebral Cortex 11(1), 85-92. https://doi.org/10.1093/cercor/11.1.85.

Nicholson, R., Karayanidis, F., Poboka, D., Heathcote, A., Michie, P.T., 2005. Electrophysiological correlates of anticipatory task-switching processes. Psychophysiology 42(5), 540-554. https://doi.org/10.1111/j.1469-8986.2005.00350.x.

Owen, A.M., Roberts, A.C., Polkey, C.E., Sahakian, B.J., Robbins, T.W., 1991. Extra-dimensional versus intra-dimensional set shifting performance following frontal lobe excisions, temporal lobe excisions or amygdalo-hippocampectomy in man. Neuropsychologia 29(10), 993-1006. https://doi.org/10.1016/0028-3932(91)90063-E.

Polich, J., 1997. EEG and ERP assessment of normal aging. Electroencephalography and Clinical Neurophysiology/Evoked Potentials Section 104(3), 244-256. https://doi.org/10.1016/S0168-5597(97)96139-6.

Richter, A., Soch, J., Kizilirmak, J.M., Fischer, L., Schütze, H., Assmann, A., Behnisch, G., Feldhoff, H., Knopf, L., Raschick, M., Schult, A., Seidenbecher, C.I., Yakupov, R., Düzel, E., Schott, B.H., 2023. Single-value scores of memory-related brain activity reflect dissociable neuropsychological and anatomical signatures of neurocognitive aging. Human Brain Mapping 44(8), 3283-3301. https://doi.org/10.1002/hbm.26281.

Smith, A.B., Taylor, E., Brammer, M., Rubia, K., 2004. Neural Correlates of Switching Set as Measured in Fast, Event-Related Functional Magnetic Resonance Imaging. Human Brain Mapping 21(4), 247-256. https://doi.org/10.1002/hbm.20007.

Stolz, C., Endres, D., Mueller, E.M., 2019. Threat-conditioned contexts modulate the late positive potential to faces—A mobile EEG/virtual reality study. Psychophysiology 56(4), e13308. https://doi.org/10.1111/psyp.13308.

Van Dinteren, R., Arns, M., Jongsma, M.L.A., Kessels, R.P.C., 2014. P300 Development across the Lifespan: A Systematic Review and Meta-Analysis. PLoS ONE 9(2), e87347. https://doi.org/10.1371/journal.pone.0087347.

West, R., Travers, S., 2008. Differential effects of aging on processes underlying task switching. Brain Cogn 68(1), 67-80. https://doi.org/10.1016/j.bandc.2008.03.001.

Zelazo, P.D., Craik, F.I.M., Booth, L., 2004. Executive function across the life span. Acta Psychologica 115(2-3), 167-183. https://doi.org/10.1016/j.actpsy.2003.12.005.

Zimmermann, P., Fimm, B., 1992. Testbatterie zur Aufmerksamkeitsprüfung:(TAP). Psytest.
